# Gene expression is the main driver of purifying selection in large penguin populations

**DOI:** 10.1101/2023.08.08.552445

**Authors:** Emiliano Trucchi, Piergiorgio Massa, Francesco Giannelli, Thibault Latrille, Flavia A. N. Fernandes, Lorena Ancona, Nils Chr Stenseth, Joan Ferrer Obiol, Josephine Paris, Giorgio Bertorelle, Céline Le Bohec

## Abstract

Purifying selection is the most pervasive type of selection, as it constantly removes deleterious mutations arising in populations, directly scaling with population size. Highly expressed genes appear to accumulate fewer deleterious mutations between divergent species’ lineages (known as E-R anticorrelation), pointing towards gene expression as an additional driver of purifying selection. However, estimates of the effect of gene expression on segregating deleterious variants in natural populations are scarce, as is an understanding of the relative contribution of population size and gene expression to purifying selection. Here, we analyse genomic and transcriptomic data from two natural populations of closely related sister species with different demographic histories, the Emperor penguin (*Aptenodytes forsteri*) and the King penguin (*A. patagonicus)*, and show that purifying selection at the population-level depends on gene expression rate, resulting in very high selection coefficients at highly expressed genes. Leveraging realistic forward simulations, we estimate that the top 10% of the most highly expressed genes in a genome experience a selection pressure corresponding to an average selection coefficient of -0.1, which decreases to a selection coefficient of -0.01 for the top 50%. Gene expression rate can be regarded as a fundamental parameter of protein evolution in natural populations, maintaining selection effective even at small population size. We suggest it could be used as a proxy for gene selection coefficients, which are notoriously difficult to derive in non-model species under real-world conditions.

## Introduction

Protein evolution is constrained by purifying selection, which prevents changes in the underlying gene sequence with a deleterious effect on organismal fitness from spreading in natural populations. The intensity of purifying selection on deleterious mutations is directly correlated with the effective size of a population (*N_e_*; Charlesworth 2009, Akashi et al 2012), determined by species-specific life history traits and population-specific demographic trajectories (Figuet et al 2016, Chen et al 2017), and with the selection coefficient (*s*) of each mutation. However, genes with a globally high expression rate across tissues show a slow rate of accumulation of deleterious substitutions (Duret and Mouchiroud 2000, Pal et al 2001, Zhang and Yang 2015), suggesting high selection coefficients on any mutation appearing in them. Such an inverse correlation between the rate of evolution and gene expression (so-called *E-R anticorrelation*) could be caused by the strong selection acting against the toxic accumulation of misfolded or mis-interacting proteins in cells (Yang et al 2012, Park et al 2013, Wu et al 2022, but see Bédard et al 2022 for more hypotheses about the causes of E-R anticorrelation). Assuming that proteins are selected for their conformational stability (*i.e.*, the protein is folded or not) or for protein– protein interaction (*i.e.*, the protein is bounded or not to other proteins), the intensity of purifying selection acting on the protein can be theoretically derived as a function of both gene expression and effective population size (Latrille & Lartillot 2021), but so far the predictions of these models have not been tested empirically in an integrated dataset.

Evidence for E-R anticorrelation has been found in several interspecific comparisons by estimating fixation rates (*d*) of nonsynonymous (*N*) over synonymous (*S*) mutations (i.e., *d_N_*/*d_S_*) in genes with different expression rates (Slotte et al 2011, Zhang and Yang 2015, Joseph et al 2017). Considering diversity at the population level, E-R anticorrelation should explain differences in nonsynonymous and synonymous segregating polymorphisms (*p*) across genes (i.e., *p_N_*/*p_S_* or as the corrected estimate *π_N_*/*π_S_*). Although such a pattern has been observed in a few wild populations (Carneiro et al 2012, Williamson et al 2014, Hodgins et al 2016, Galtier et al 2016), recent laboratory experiments on model organisms have instead provided contrasting results (Wu et al 2022, Shibai et al 2022). More importantly, the relative contribution of gene expression and effective population size to purifying selection has not been empirically explored. Theory predicts that the efficiency of purifying selection depends on the product of effective population size and selection coefficient to be much larger than 1. We can therefore ask whether genes with high expression levels are characterised by large enough selection coefficients so that purifying selection still exerts its effect even when populations are small. On the other hand, understanding the range of selection coefficient values across genes would help identify those genes which are more vulnerable to decreasing population size.

Here, we use two natural populations of closely related sister species, the Emperor and the King penguins (*Aptenodytes forsteri* and *A. patagonicus*), with different demographic histories (Trucchi et al 2014, Cristofari et al 2016, 2018), to test the following hypotheses. First, if the selection coefficient of a gene is mainly determined by its expression rate, we should observe a decline in the effect of purifying selection (e.g. *π_N_*/*π_S_*) with increasing expression rate and such decline should be determined by a corresponding decline in missense polymorphism only. Our second question concerns the relative weight of population size (*N_e_*) and gene expression (*s*) in driving purifying selection. When comparing populations of different sizes, smaller populations show lower diversity at both neutral and deleterious sites, but higher *π_N_*/*π_S_* because of larger drift which reduces the efficacy of purifying selection. If population size is the main driver of purifying selection (1/*N_e_* > *s* across the whole range of gene expression), we expect that both the diversity and the *π_N_*/*π_S_* differences between the two populations of different sizes will be the same across the whole range of gene expression. Conversely, if high gene expression is the main driver of purifying selection (*s* > 1/*N_e_* for highly expressed genes), we expect the difference in diversity between the two populations of different size to decline with increasing gene expression rate for deleterious sites but not for neutral ones. Finally, we use realistic forward simulations of evolving populations to estimate the range of selection coefficients producing the same effects of purifying selection as observed in natural populations of Emperor and King penguins.

## Results and Discussion

We use high-coverage whole-genome data of 24 individuals per species to estimate patterns of genetic diversity, and whole transcriptome data of five tissues from three young individuals per species to estimate global mRNA expression levels. Young age class was chosen for this study as genes broadly expressed in early life stages have been shown to be the most affected by purifying selection (Cheng and Kirkpatrick 2021). Both Emperor and King penguins feature single, large and quasi-panmictic populations (Cristofari et al 2016, 2018), but they show different levels of genetic diversity (Fig. 1A), corresponding to their different ecological adaptations and past demographic dynamics (Cristofari et al 2016, 2018, Cole et al 2022). As a consequence of the historically larger effective population size in the Emperor penguin, this species has a higher proportion of segregating variants, a lower proportion of fixed derived variants, and a lower proportion of segregating nonsynonymous over synonymous variants (Fig. 1B); however, the two sister species show a minor difference in the proportion of fixed nonsynonymous over synonymous differences, given their relatively short time since species divergence. Both gene-by-gene estimates of diversity (nucleotide diversity: *π*) and expression rate (normalised as transcripts per million, TPM) are highly correlated between the two species (Fig. 1C, D), thus minimising any confounding effect of sequence and expression divergence in our downstream analyses.

**Figure 1.**
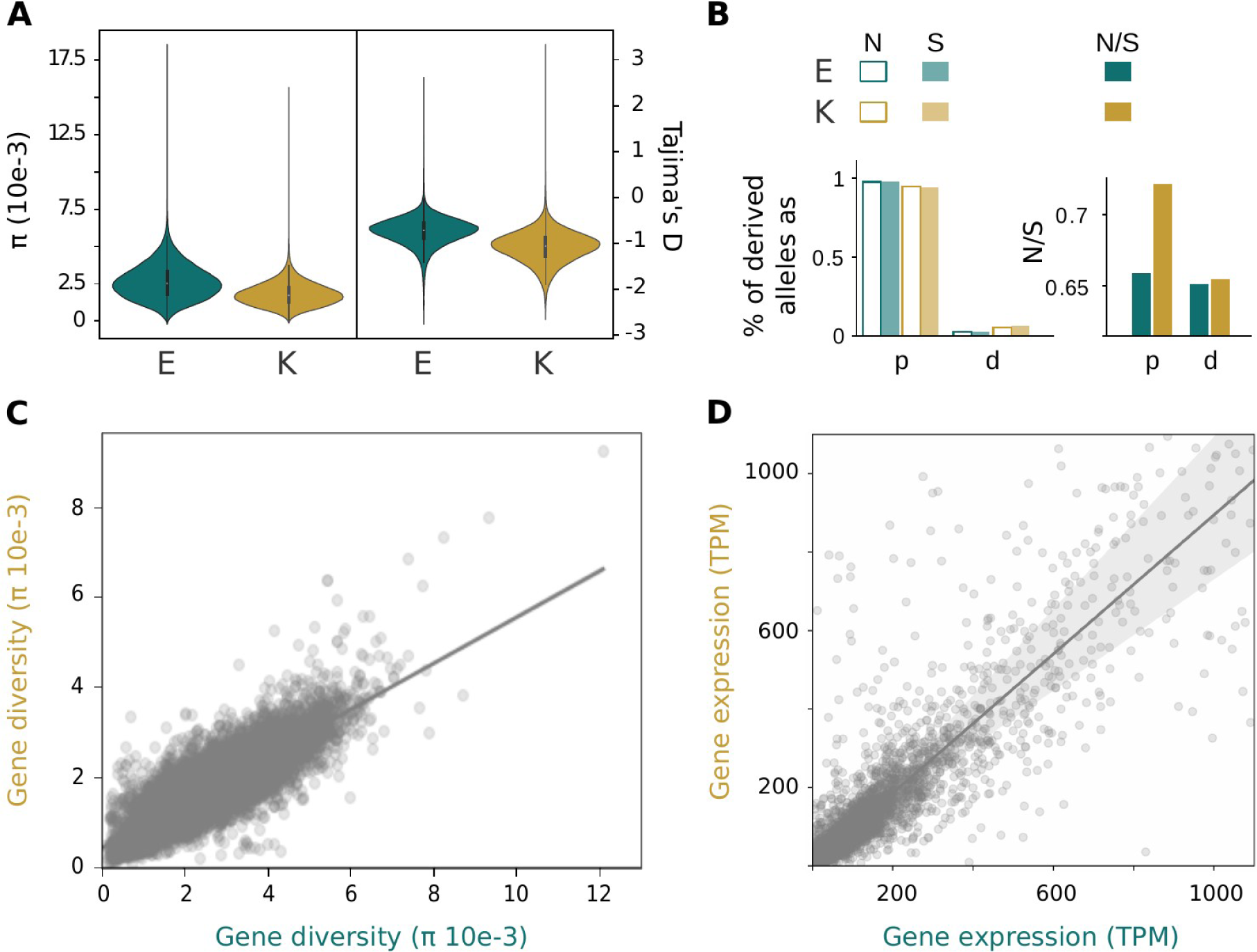
Patterns of genetic diversity and gene expression in Emperor (E, teal) and King (K, gold) penguins. **A.** Distribution of nucleotide diversity (*π*) and Tajima’s D in 50 kb genomic windows; **B.** Proportion of derived alleles as segregating variants (*p*) or fixed differences (*d*) at synonymous and nonsynonymous sites (left panel) and estimates of *pN/pS* and *dN/dS* (right panel); **C**. Per gene comparison of nucleotide diversity between King and Emperor penguins; **D.** Per gene comparison of expression rate between King and Emperor penguins, quantified as transcripts per million (up to TPM = 1100; see Supp. Fig. 5 for the whole expression range).

### Purifying selection more efficiently removes nonsynonymous segregating variants in genes while expression rate increases

Corrected estimate of purifying selection on segregating variants per gene, *π_N_*/*π_S_*, clearly declines with increasing gene expression rate (Fig. 2A), dropping by 70-80% across the whole range of gene expression in both species. These results hold regardless of binning or not the genes in percentiles of expression rate (Supp. Fig. 6) and are consistent with the E - R anticorrelation found in several taxa at the interspecific divergence level as shown in Zhang and Yang (2015) (Supp. Fig. 7). As expected, also the rate of fixation of nonsynonymous over synonymous mutations (*d_N_*/*d_S_*) declines with gene expression rate in both species, even if divergence estimates are null for many genes given the shallow split time between two penguin species (Supp. Fig. 6, 7). E-R anticorrelation appears also if we analyse the expression rate of segregating sites across all genes together (mRNA sequencing coverage per site normalised as count per million reads - CPM) in order to take into account heterogeneous expression rate among exons: again, counts of nonsynonymous over synonymous variants in bins of 0.05 CPM, from 0 to 5 CPM, are inversely correlated with expression rate (Supp. Fig. 9). The decline of *π_N_*/*π_S_* with increasing gene expression rate is due to the decreasing count of nonsynonymous variants in highly expressed genes, whereas the count of synonymous variants is stable across the whole gene expression range in both species (Fig. 2B, C). More importantly, the difference in the counts of synonymous variants between the two penguin populations is also stable and always significant (Kolmogorov-Smirnov test *p*-value << 0.005), whereas the difference in the counts of nonsynonymous variants decrease with increasing gene expression, with this difference disappearing in the upper 50-60% of gene expression rate (Fig. 2C). This result supports the hypothesis that gene expression is a major driver of purifying selection for highly expressed genes, which are then expected to show very large selection coefficients (*s* > 1/*N_e_*). As theoretically predicted (Latrille & Lartillot 2021), the rate of purifying selection appears to linearly decrease with the logarithm of the expression rate (Fig. 2A). After estimating the change in rate of purifying selection (*π_N_*/*π_S_*) as a function of the effective population size of the two penguin species in log scale (Supp. Fig. 8), we show that all estimated slopes are statistically different from zero and negative. However, the slope estimates are not significantly different from each other and their confidence intervals overlap (Supp. Fig. 8). Compatible with the assumptions that proteins are selected for their conformational stability or for protein–protein interaction, these results suggest that both the effects of effective population size and gene expression can be considered together in integrated models of evolution. However, they should be assessed more thoroughly, by comparing more population sizes. On a different note, even if gene expression has been suggested to be one of the causes of non-neutrality in synonymous variants in yeast (Shen et al 2022), or that codon usage bias is more intense in highly expressed genes (Frumkin et al 2018), gene expression rate does not appear to perturb synonymous variation in our datasets from two vertebrate species (Fig. 2C).

**Figure 2.**
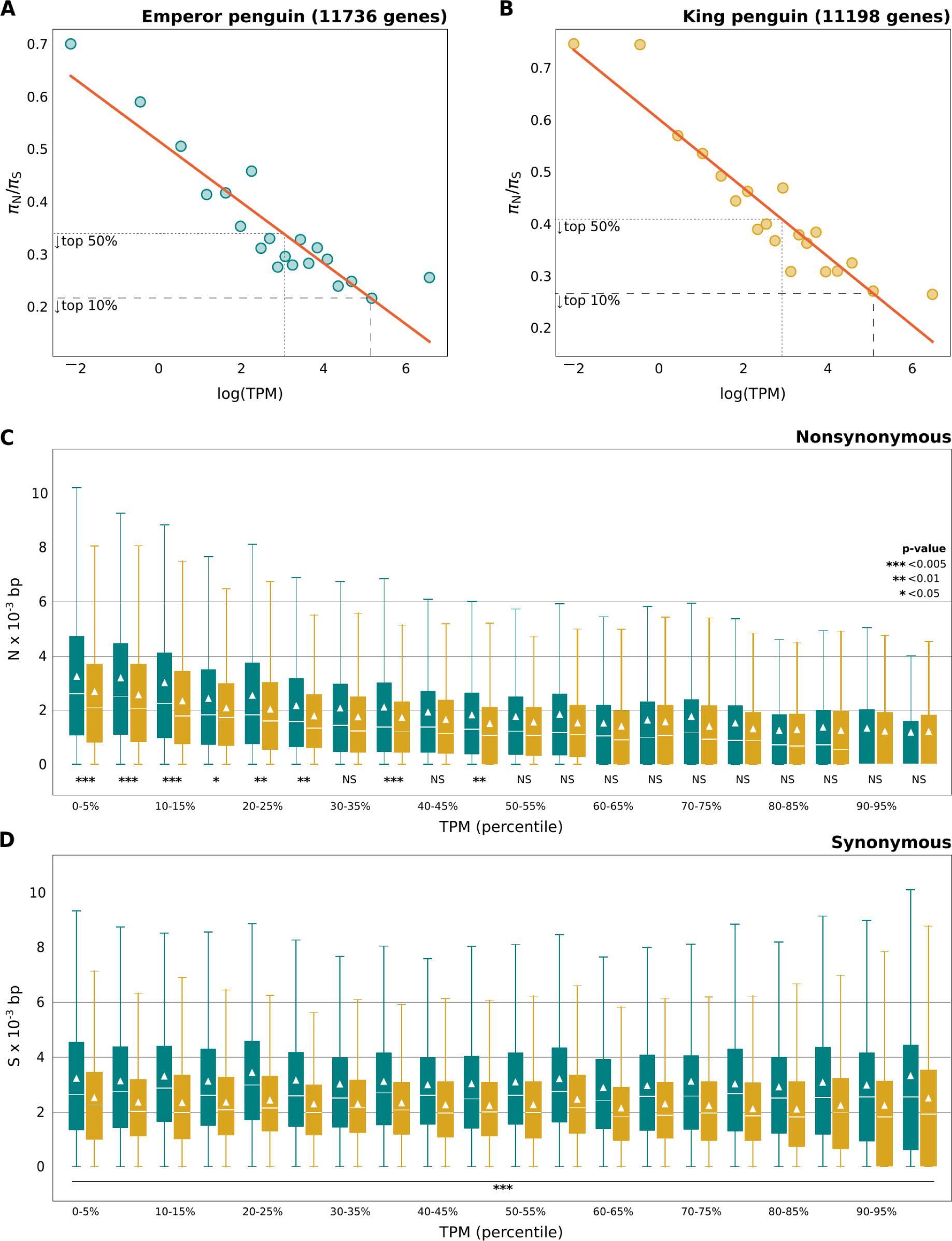
Increasing purifying selection with gene expression in Emperor (teal) and King (gold) penguins. Estimates of *πN*/*πS* (**A, B**), and average number of nonsynonymous (**C**) and synonymous (**D**) segregating variants (normalised per 1000 bp of coding sequence) in genes binned by 5% percentiles of expression rate (normalised as TPM). Slope of the linear regression (Emperor penguin: *χ^E^* = -0.058, *R^2^ = 0.848;* King penguin: *χ^K^* = 0.066, *R^2^* = 0.877) is shown as a solid red line and values of *πN*/*πS* for the top 50% (small dashes) and 10% (large dashes) of the most highly expressed genes are indicated by a dashed grey line in panels A and B. Median (solid white line) and mean (white triangle) is shown in each boxplot in panels C and D. Statistical significance for the difference in the distribution of synonymous and nonsynonymous variants per percentile between the two species is shown in panels B and C.

### Purifying selection more efficiently prevents nonsynonymous segregating variants from increasing in frequency in genes with higher expression rate

The derived allele frequency spectrum of nonsynonymous variants with expression rate higher than 0.3 CPM is depleted in medium-high frequency categories, while there is no difference in the derived allele frequency spectrum of synonymous variants across the whole expression range (Fig. 3). Changing the arbitrary threshold to discriminate between low and high expression rate, or using more than two categories of expression rate (low: < 0.3 CPM, medium: 0.3-2 CPM, high: > 2 CPM) does not change the observed pattern (Supp. Fig. 10). The pattern holds when all nonsynonymous and synonymous variants are used in the allele frequency spectrum estimate (Supp. Fig. 10) as well as when one nonsynonymous and one synonymous variant are randomly sampled from each gene (Fig. 3), thus excluding the possibility that few genes with many variants (*i.e.,* pseudoreplication) drive our observation.

**Figure 3.**
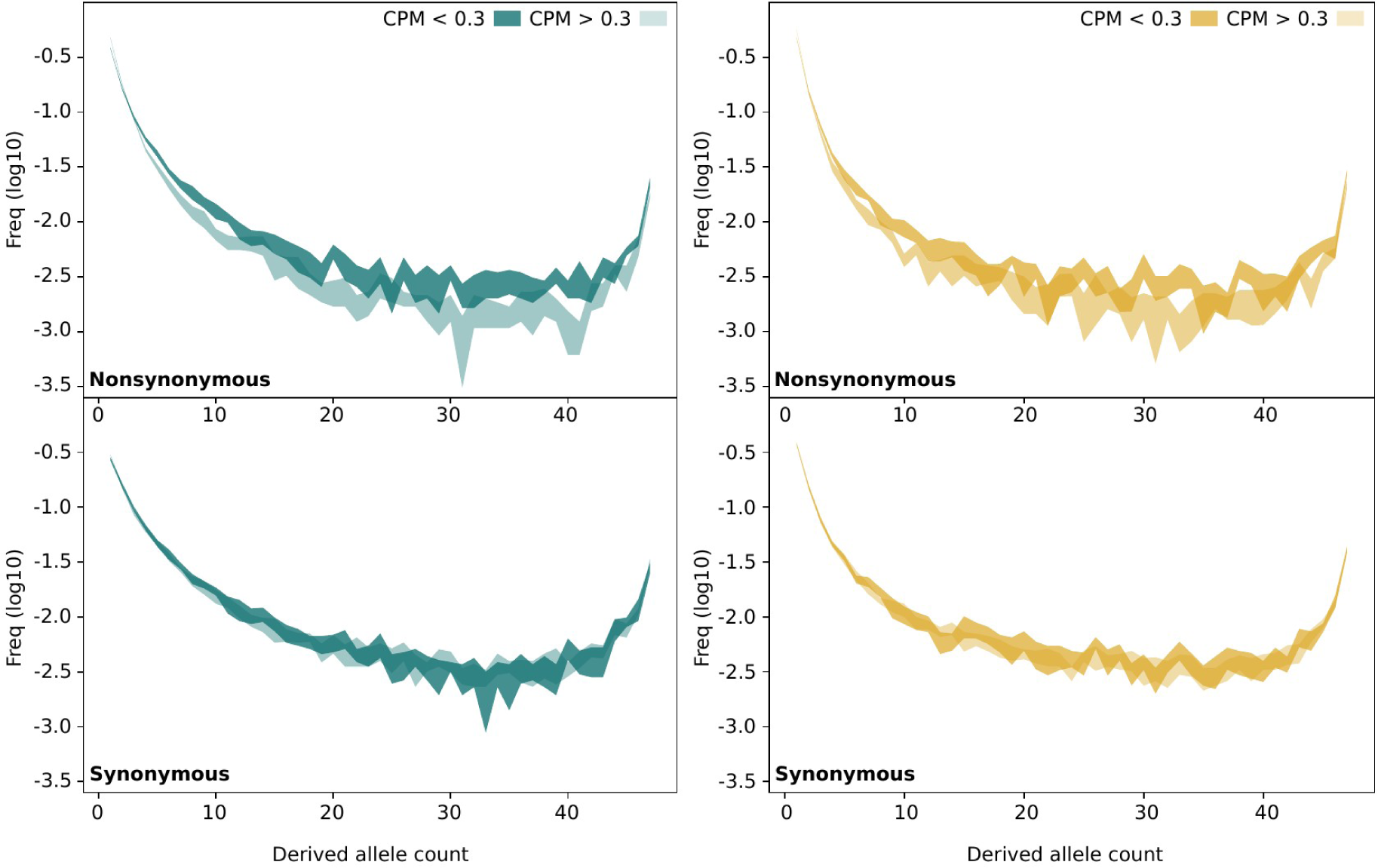
Nonsynonymous variants in highly expressed genes segregate at lower frequency in Emperor (teal) and King (gold) penguins. Site frequency spectra of ten random resampling (95% distribution) of one nonsynonymous (upper panels) and one synonymous (lower panels) variant per gene with lower (dark shade) or higher (light shade) than 0.3 CPM mRNA expression. The relative frequency of each count class is log10 transformed. Of note, nonsynonymous variants in highly expressed genes show a higher frequency at low derived allele count classes than in lowly expressed genes.

### Purifying selection in the top 10% of highly expressed genes largely exceeds the effect of 100,000-individuals effective population size

In simulated populations, under either Wright-Fisher or more realistic non Wright-Fisher models, median *p_N_*/*p_S_* (same as *π_N_*/*π_S_* when using simulated data) across genes declines from 1.8 to 0.9, while population size increases from 1,000 to 100,000 individuals (Fig. 4). Such values of *p_N_*/*p_S_* are much higher than the values observed in penguin populations for genes in the top 50% or top 10% of expression rate (Fig. 2A, Supp. Fig. 6). In these models, the effect of population size on purifying selection was explored by simulating a set of realistic values for mutation and recombination rate, synonymous to nonsynonymous ratio, selection and dominance coefficient distributions, coding sequence length and gene numbers. In particular, new mutations were given a selection and dominance coefficient (*h*-mix) based on a nearly neutral prior distribution (Kim et al 2017, Kyriazis et al 2021), meaning that most of the mutations are weakly deleterious. To reproduce *p_N_*/*p_S_* values as those observed in highly expressed genes in both penguin species, we designed a more extreme selection scenario: all nonsynonymous mutations appearing in a gene were given a fixed selection coefficient of -0.1, -0.01 or -0.001 (100 replicated genes per selection coefficient) and a dominance coefficient derived from the *hs* relationship (Henn et al 2016). In realistic non Wright-Fisher models with a selection coefficient of -0.01, *p_N_*/*p_S_* decreases below 0.4 (Fig. 4) while a selection coefficient of -0.1 results in *p_N_*/*p_S_* below 0.3, as largely observed in genes in the top 50% and 10% of expression rate, respectively (Fig. 2A, Supp. Fig. 6). Such high selection coefficients are then expected to be effective even when the population size is small (*i.e.*, *s* >> 1/*N_e_*, per *N* = 1,000), thus buffering the effects of changing population size as it was also suggested for the vast majority of X-linked genes in *Drosophila* (Andolfatto et al 2011). However, we also observe more variance in simulations with smaller population sizes (Supp. Fig. 11).

**Figure 4.**
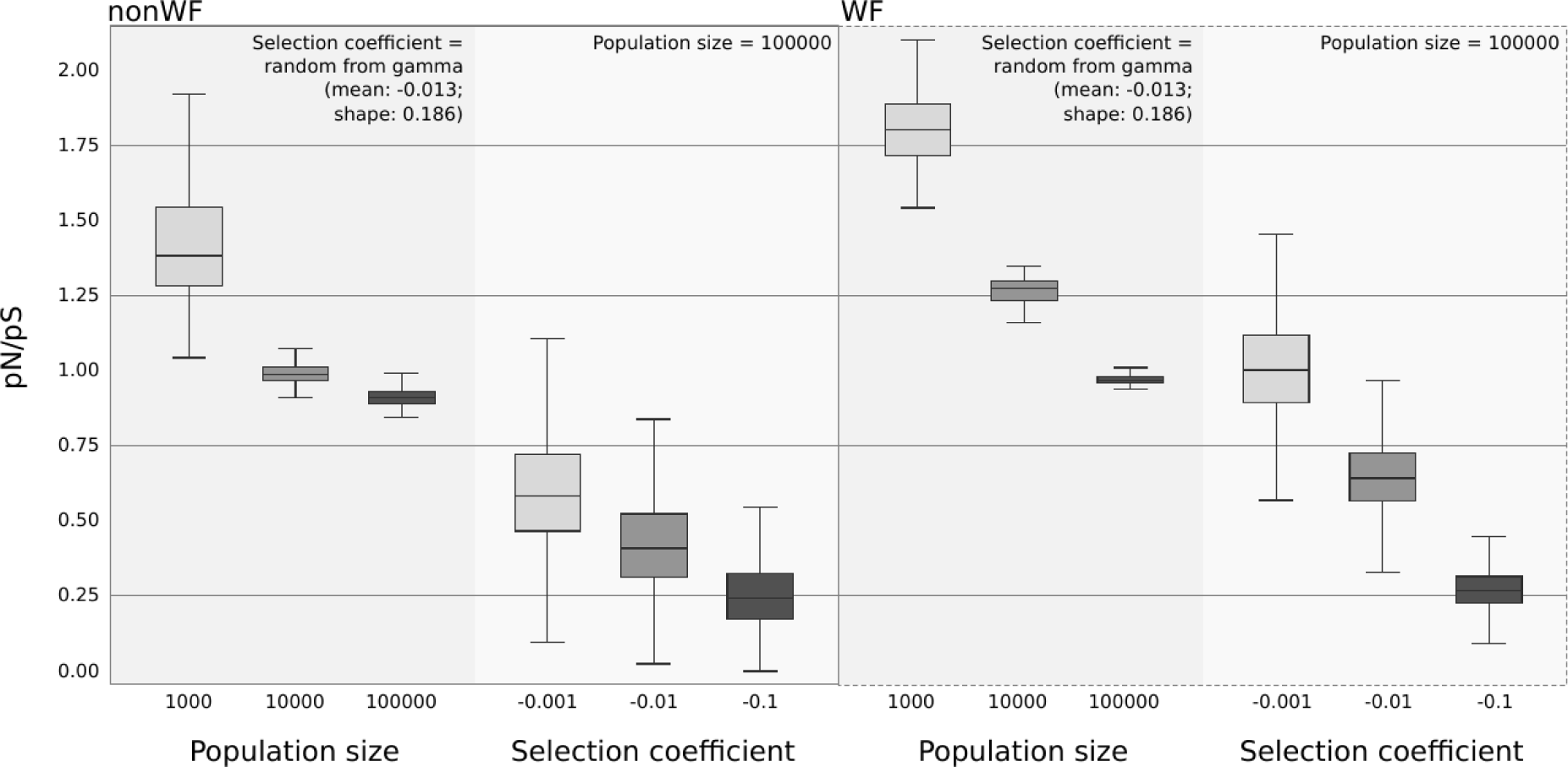
Population size and gene-specific extreme selection coefficient explain low observed *pN/pS* values in simulations. Distribution of *pN*/*pS* (when using simulated genomic data *pN*/*pS* is the same as *πN*/*πS*) across 1000 genes simulated under nonWrightFisher (nonWF, left, solid border) and WrightFisher (WF, right, dashed border) models with effective population size from 1,000 to 100,000 (darker grey background) and across 100 genes with selection coefficient from -0.001 to -0.1 (lighter grey background). Note that the dominance coefficient is set according to the *h-mix* or *hs* models in simulations testing different population sizes or selection coefficients, respectively. More efficient purifying selection in nonWF models, where effective population size tends to be lower than in WF models, can be explained by the fact that, in such models, individuals with high fitness can survive and reproduce for multiple generations (Haller and Messer 2019).

### Gene expression can be used as a proxy of the distribution of gene selection coefficients in natural populations of non-model species

Variants with highly deleterious effects on individual fitness are expected to be immediately lost in natural populations. Consistent with this expectation, the highly deleterious variants (less than1000 HIGH effect SNPs per species) predicted by SNPeff (Cingolani et al 2012) in each penguin population show a much lower average expression level than weakly deleterious (MODERATE effect) and nearly-neutral (LOW effect and synonymous) variants (Tab. 1). This observation means that HIGH effect variants are mainly present in lowly expressed genes with limited impact on fitness. Expression level is even lower in the very few fixed differences with HIGH effect (Tab. 1), thus supporting our hypothesis. As site-specific expression of highly deleterious variants (mainly start/stop codons loss/gain and splice acceptor/donor variants) could be biassed in mature mRNA sequencing, we also estimated the expression of highly deleterious variants as the expression of the gene they belong to. Even applying a rather conservative test, the expression of genes with predicted highly deleterious variants is on average three times lower than all genes (Kolmogorov-Smirnov test *p-*value = 0.00095) in the King penguin and slightly lower, even if not significant, in the Emperor penguin. As previously suggested for the distribution of dominance coefficients in a model plant species (Huber et al 2018), gene expression should be taken into account when using predictions of fitness effects and, more generally, when using such predictions to calculate the genetic load in populations of conservation concern (Bertorelle et al 2022). In fact, predicted highly deleterious variants could be on lowly expressed genes, thus with little contribution to individual or population fitness.

**Table 1.** Expression rate by predicted fitness effect. LOWsyn: low effect and synonymous; MDR: moderate effect; HIGH: high effect.

| PREDICTED FITNESS EFFECT | LOWsyn | LOWsyn | MDR | MDR | HIGH | HIGH |
| --- | --- | --- | --- | --- | --- | --- |
| Species | Emperor | King | Emperor | King | Emperor | King |
| Total variants | 73501 | 57088 | 47183 | 41352 | 934 | 840 |
| →Average (stdev) Z-normalised CPM | 1.5 (5.02) | 1.3 (4.56) | 0.78 (2.85) | 0.78 (2.86) | 0.24 (1.97) | 0.14 (0.88) |
| Fixed differences | 1846 | 3500 | 1166 | 2229 | 16 | 44 |
| →Average (stdev) Z-normalised CPM | 1.41 (4.12) | 1.22 (4.14) | 1.4 (6.31) | 0.93 (3.39) | 0.08 (0.54) | 0.02 (0.57) |
| Average count of derived alleles in segregating variants | 6.17 | 6.25 | 4.75 | 4.73 | 5.14 | 5.48 |

### Concluding remarks

Overall, our study provides evidence that gene expression is a fundamental driver of purifying selection in natural populations and to a higher extent than population size for highly expressed genes. About half of the genes in a genome, which are likely responsible for basic cellular and molecular functions (Boyle et al 2017), are under a strong selective constraint preventing deleterious sequence changes even when population size declines to about 1,000 individuals. Selection coefficients on the top 10% of the expressed genes could be so high to buffer even smaller population size (ca. 100 individuals). Below this order of magnitude, random effects would necessarily prevail in the proteins’ evolutionary trajectory. Importantly, gene expression can be used as a proxy of the gene selection coefficient, which is notoriously difficult to study in natural populations of non-model species (Huber et al 2017). Gene expression data are easier to collect than selection coefficients and are usually highly conserved across closely related species (Fig. 1D), so that they can be used to refine estimates of genetic load (Bertorelle et al 2022) in natural populations of conservation concern.

### Data availability

Genomic and transcriptomic raw reads are publicly available at ENA database with Project accession number XXX (genomic raw reads to be submitted) and PRJEB64484, respectively.

The filtered SNPs dataset is available here: 10.5281/zenodo.10688854. Bioinformatic scripts are available here: github.com/emitruc/ExpressionLoad; github.com/ThibaultLatrille/PenguinExpression; github.com/PiergiorgioMassa/penguin_gene_expression_slimulations.

## Supporting information

Supplementary File 1, Samples sheet

## Acknowledgements

We thank Fabrizio Mafessoni, Paolo Gratton and Robin Cristofari for helpful comments and suggestions on the analyses, and Federica Pirri for RNA data production. ET was supported by PNRA_16 00164 (“Programma Nazionale di Ricerca in Antartide”. Bando PNRA 5 aprile 2016, n. 651. – Linea B “Genomica degli adattamenti estremi alla vita in Antartide”). The study was supported by the Institut Polaire Français Paul-Emile Victor (IPEV) within the framework of the Program 137-ANTAVIA (PI: CLB), by the Centre Scientifique de Monaco with additional support from the LIA-647 and RTPI-NUTRESS (CSM/CNRS-UNISTRA), by the Centre National de la Recherche Scientifique (CNRS) through the Programme Zone Atelier de Recherches sur l’Environnement Antarctique et Subantarctique (ZATA). This study was approved by the French ethics committee (last: APAFIS#29338-2020070210516365) and the French Polar Environmental Committee, and permits to handle animals and access breeding sites were delivered by the “Terres Australes et Antarctiques Françaises” (TAAF). The sequencing service was provided by the Norwegian Sequencing Centre (www.sequencing.uio.no), a national technology platform hosted by the University of Oslo and supported by the “Functional Genomics” and “Infrastructure” programs of the Research Council of Norway and the Southeastern Norway Regional Health Authorities. Bioinformatic analyses were performed on the HPC clusters of the Department of Life and Environmental Sciences (“HappyComputing@DiSVA”), Marche Polytechnic University, and of the Department of Life Sciences and Biotechnology, University of Ferrara.

## Author contributions

ET designed the study and secured fundings together with GB, extracted the DNA samples, analysed the genomic and transcriptomic data, and wrote the manuscript; PM and FG built, performed and analysed the genomic simulations, and wrote the relevant section in the Methods; TL performed the regression analyses between purifying selection, expression rate and population size, and wrote the relevant text (section 3) in the Extended Methods; NCS contributed to the pilot study leading to this work; MB, CC collected the samples for RNA analyses and contributed to the manuscript; FANF, LA, JFO, JP and GB discussed the results and contributed to the manuscript; CLB coordinated the project logistics and the samples collection and the associated fundings, discussed the results and contributed to the manuscript. All authors contributed to the development of the paper.

## Extended methods and supplementary information

### Genomic data

#### 1.1 Samples collection and storage

Genomic DNA extractions of 24 King penguins (*Aptenodytes patagonicus* from South Georgia, Crozet archipelago, and Heard island), and 24 Emperor penguins (*Aptenodytes forsteri* from Terre Adélie and Dronning Maud Land, Antarctica), were selected from samples used in Cristofari et al (2016) and Cristofari et al (2018). In addition, four Gentoo penguin (*Pygoscelis papua* from Crozet archipelago) and two Adelie penguins (*Pygoscelis adeliae* from Terre Adélie, Antarctica, 2007) were collected during the field campaigns of the French IPEV programme 137 in 2017 and 2007, respectively (Supp. File 1). Samples were stored in ETOH (muscle biopsy) or in Queen Lysis buffer (blood samples), frozen at -80°C from the field to the lab.

#### 1.2 DNA extraction, pooled library preparation and sequencing

DNA extraction was performed using DNeasy Blood and Tissue kit (Qiagen) following manufacturer’s instructions. Whole genome sequencing libraries were prepared and sequenced at the Norwegian Sequencing Centre, Oslo, using Illumina pcr-free single or dual-indexing kits. To minimise batch effects, genomic samples from different species and different localities were randomised in six libraries and sequenced on 1-3 lanes of Illumina HiSeq2500 and HiSeq4000 aiming at 20X coverage depth. Raw reads are publicly available at ENA database (XXX, to be made available).

#### 1.3 Variant calling, filtering and annotation

After Illumina adapters trimming and quality filtering with Trimmomatic (Bolger et al 2014) and URQT (Modolo and Lerat 2015), respectively, fastq reads were mapped to the *Aptenodytes forsteri* reference genome (ASM69914v1; RefSeq assembly accession: GCF_000699145.1) using *bwa mem* (v0.7.15; Li 2013), converted to bam files and sorted with *samtools* (v0.1.19; Li et al 2009), keeping only reads with phred-scaled mapping quality higher than 10. Duplicated reads were removed with *picard-tools* (v1.98) and bam files were assigned to individual samples by adding ID read groups with picard-tools.

Small variants (SNPs, indel and MNPs) were called with *freebayes* (Garrison and Marth 2012) using reference genome scaffolds longer than 100Kb (481 in total), all samples grouped per species (-*-populations* flag), and a minimum phred-scaled mapping quality of 20. Resulting vcf files (one per scaffold) were then filtered: *i)* MNPs were first broken down into SNPs using *vcfallelicprimitives* script in *vcflib* (with -k -g flags; Garrison et al 2022); *ii)* variants were then filtered for quality (QUAL > 30), strand bias (SAF > 0 & SAR > 0), read placement bias (RPL > 0 & RPR > 0), and type of variants (TYPE = snp) with *vcffilter* script in vcflib; *iii)* SNPs were finally filtered using vcftools (v0.1.15; Danecek et al 2011) for minimum coverage depth of 3 reads per individual (--minDP 3; individual genotypes discarded if below threshold), mean maximum coverage depth of 50 (--max-meanDP 50 which is ca. three times the average individual coverage depth; whole locus discarded if above threshold), and retained only if biallelic across all samples. A total of 44 sex-linked scaffolds were identified and removed from the dataset by running samtools *idxstats* on all bam files and the SATC (Nursyifa et al 2022) Rscript on the resulting data. SNPs were annotated using SNPeff (Cingolani et al 2012) and the GCF_000699145.1 genome annotation. Annotated vcf files are available upon request. Mean coverage depth of filtered SNPs per allele was 6.93X, standard deviation 1.07X.

#### 1.4 SNPs polarisation in ancestral and derived alleles

Annotated vcf files were parsed using a custom python script (*vcf2missenseFreq.2d.py;* https://github.com/emitruc/ExpressionLoad) to decide on the derived allele. Using the Emperor and King penguins samples as ingroups and the Adelie and Gentoo penguins samples as outgroups, we defined all of the possible configurations of a globally polymorphic site (Supp. Fig. 1). After assessing the most likely ancestral allele based on our algorithm (Supp. Fig. 1), for each SNP position, we calculated the joint derived allele counts for King and Emperor penguin samples and summarised the data in the table *daf.joint* (Supp. Tab. 1) including the following information (column labels are in brackets):

- counts of derived alleles and total alleles in Emperor penguin (*der1*; *tot1*), King penguin (*der2*; *tot2*), and Adélie and Gentoo penguins (*der_out*; *tot_out*) samples;
- average allele coverage (*avgCov*);
- ancestral (*ref*) and derived (*alt*) allele;
- genomic site type (*vartype*) based on the first annotation by SNPeff as *missense* if containing the word ‘missense’, *synonymous* if containing the word ‘synonymous’, *intergenic* if containing the word ‘intergenic’, *intronic* if containing the word ‘intron’, *else* otherwise;
- genomic site predicted effect (*effect*) based on the the first annotation description as HIGH, MODERATE, LOW, MODIFIER;
- a flag whether the polymorphic site was originally called as MNP or SNP by freebayes (*flagQual*: *haplo*, *snp*);
- a flag to track how the derived allele was called (*flagPol*) on the basis of the options shown in Supplementary Figure 1.

**Supplementary Table 1.** Example of SNPs recorded in the summary table *daf.joint*.

| scaffold | position | flagPol | flagQual | der1 | tot1 | der2 | tot2 | der_out | tot_out | avgCov | ref | alt | vartype | effect |
| --- | --- | --- | --- | --- | --- | --- | --- | --- | --- | --- | --- | --- | --- | --- |
| NW_008793941.1 | 85516 | InFixAnc | snp | 0 | 48 | 0 | 44 | 6 | 8 | 7.2 | G | A | intron | MODIFIER |
| NW_008793941.1 | 85528 | unfolded | snp | 3 | 48 | 46 | 46 | 0 | 8 | 6.6 | G | A | intron | LOW |
| NW_008793941.1 | 85558 | allFix | snp | 0 | 48 | 0 | 46 | 8 | 8 | 6.4 | T | C | synonymous | LOW |
| NW_008793941.1 | 85657 | unfolded | snp | 0 | 48 | 1 | 48 | 0 | 12 | 6 | C | T | synonymous | LOW |
| NW_008793941.1 | 85734 | unfolded | snp | 2 | 48 | 0 | 48 | 0 | 12 | 6.9 | C | T | intron | MODIFIER |
| NW_008793941.1 | 85746 | allFix | snp | 0 | 48 | 0 | 48 | 12 | 12 | 6.8 | G | A | intron | MODIFIER |
| NW_008793941.1 | 85762 | unfolded | snp | 5 | 48 | 0 | 48 | 0 | 10 | 6.6 | C | T | intron | MODIFIER |
| NW_008793941.1 | 85788 | InFixAnc | snp | 0 | 48 | 0 | 46 | 4 | 8 | 7.1 | A | C | intron | MODIFIER |
| NW_008793941.1 | 85790 | unfolded | snp | 0 | 48 | 15 | 46 | 0 | 8 | 7.1 | T | C | intron | MODIFIER |
| NW_008793941.1 | 85800 | unfolded | snp | 2 | 48 | 0 | 48 | 0 | 8 | 6.9 | C | T | intron | MODIFIER |
| NW_008793941.1 | 85804 | unfolded | snp | 1 | 48 | 0 | 48 | 0 | 8 | 6.9 | C | T | intron | MODIFIER |
| NW_008793941.1 | 85829 | unfolded | snp | 0 | 48 | 11 | 48 | 0 | 8 | 6.7 | T | C | intron | MODIFIER |

**Supplementary Figure 1.**
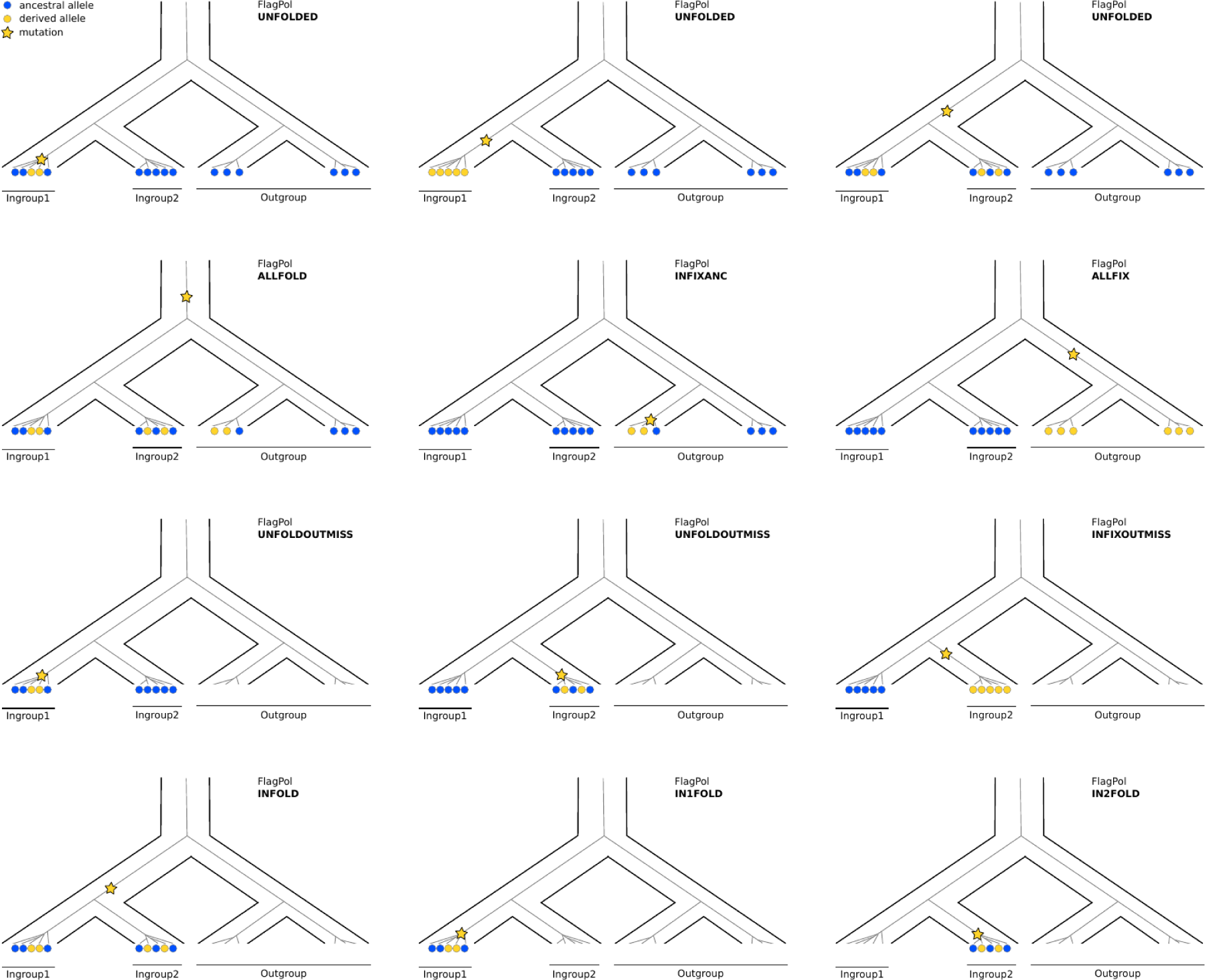
Schematic representations of the potential distribution of ancestral and derived alleles at a biallelic site across two ingroup and one outgroup samples. Ingroup1: Emperor penguin; ingroup2: King penguin; outgroup: *Pygoscelis* (Adélie + Gentoo) penguins. Blue and yellow circles represent copies of the ancestral and derived alleles. The number of allele copies per sample is only given as an example. Missing data are represented as no circles. The yellow star represents the branch and timing the mutation to the derived allele likely occurred at. For ALLFIX and INFIXOUTMISS configurations, the allele of ingroup1 is arbitrarily considered as ancestral. Ingroup1 or ingroup2 missing data configurations are not shown. This algorithm is implemented in the python script *vcf2missenseFreq.2d.py (available at* https://github.com/emitruc/ExpressionLoad).

#### 1.5 Final data filtering and sanity checks

After removing monomorphic sites across Emperor and King penguin samples (*e.g.*, INFIXANC and ALLFIX configurations in Supp. Fig. 1) from the dataset (*daf.joint.no00*, https://zenodo.org/doi/10.5281/zenodo.10688853), we checked whether the derived alleles distribution was in line with basic expectations from population genetics. To avoid downstream normalisation, we selected only sites without missing data in the target species. In addition, we choose sites which were properly polarised (flagPol=UNFOLDED) based on at least four alleles present in the outgroup and with coverage depth range per allele between 6X and 8X, *i.e.*, one standard deviation (*ca.* 1X) from the mean (*ca.* 7X). Such a narrow coverage range was applied to mitigate as much as possible the inclusion of sex-chromosome related regions, which could have been incorrectly assembled within autosomal scaffolds, and multiple-copy regions, which were not properly assembled in the reference genome sequence (see Supp. Fig. 2 for a comparison at different coverage ranges). Both types of regions are also characterised by marked deviation from Hardy-Weinberg equilibrium (HWE). Applying a coverage threshold as a filter, we could retain sites which are not in HWE due to other processes (*e.g.*, population structure, selection).

Using all intergenic sites with no missing data for both species, we summarised the joint derived allele frequency (DAF) distribution (Supp. Fig. 2). As expected, a very small proportion (note the log scale of the heatmap in Supp. Fig. 2) of derived alleles appear as segregating in both species, with a minimal covariance in the low and in the very high frequency classes. The latter suggests some mis-polarisation due to sites hit by multiple mutations (see below). Nevertheless, when setting the coverage range between 6X and 8X, co-segregating derived allele frequencies largely appear as uncorrelated between the two species as expected in case of incompletely sorted ancient variation.

**Supplementary Figure 2.**
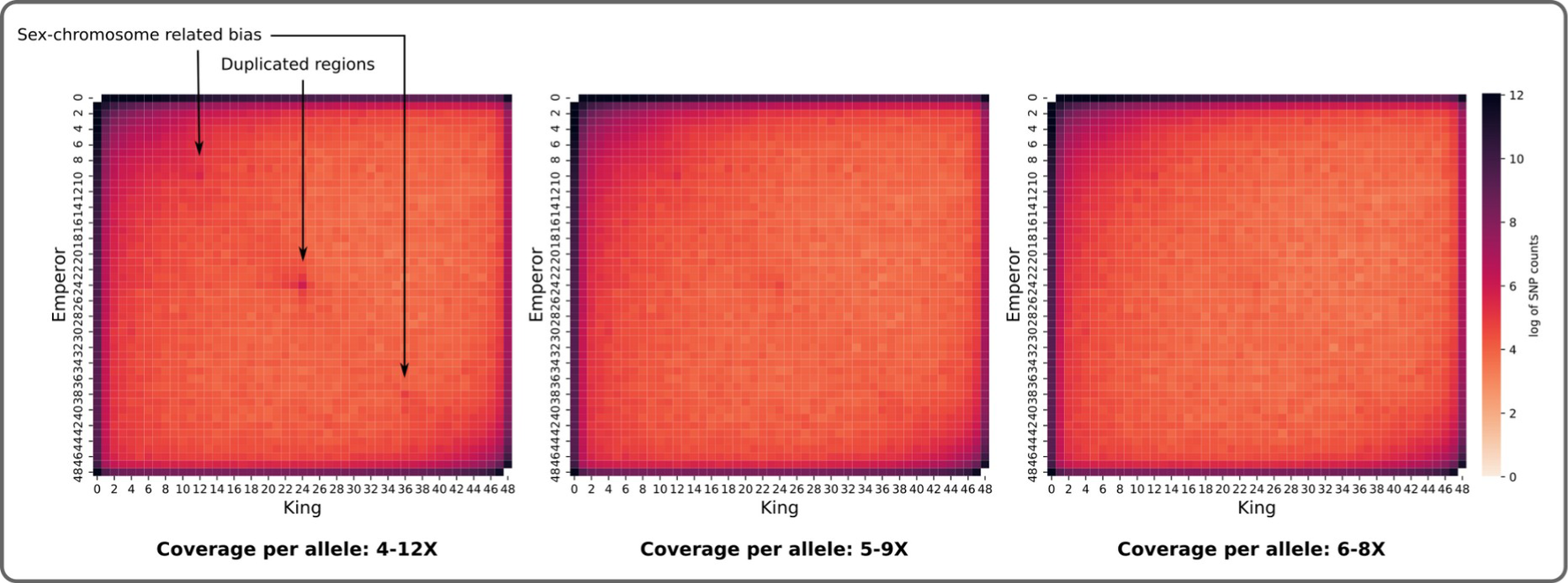
Joint derived allele frequency distributions in King and Emperor penguins samples at different ranges of coverage.

DAF were separately summarised according to the predicted annotation as intergenic, intronic, synonymous and nonsynonymous in each species (Supp. Fig. 3). In general, the shape of observed DAF is consistent with the expectations from population genetics theory. The slight (note the log scale on the y-axis in Supp. Fig. 3) increase in very high frequency variants has been commonly observed (also in human population data; Marchi and Excoffier 2020) and it is likely due to mis-polarisation of ancestral/derived alleles at sites hit by multiple mutations (Hernandez et al 2007), rather than to migration from a “ghost” population (Marchi and Excoffier 2020). DAF from intergenic sites represent the best approximation to neutrality, with a shape depending on past population demography only. On the contrary, DAF from missense sites are expected to be enriched in low frequency and depleted in medium-high frequency classes as deleterious alleles are less likely to increase in frequency in the population due to negative selection. Such a pattern is clearly appearing in both Emperor and King penguins. Synonymous sites DAF show small deviations from neutrality, likely due to linked selection within missense sites in exons. On the other hand, DAF from intronic and intergenic sites are fully overlapping with each other, suggesting very limited linked selection spanning from exons to introns.

**Supplementary Figure 3.**
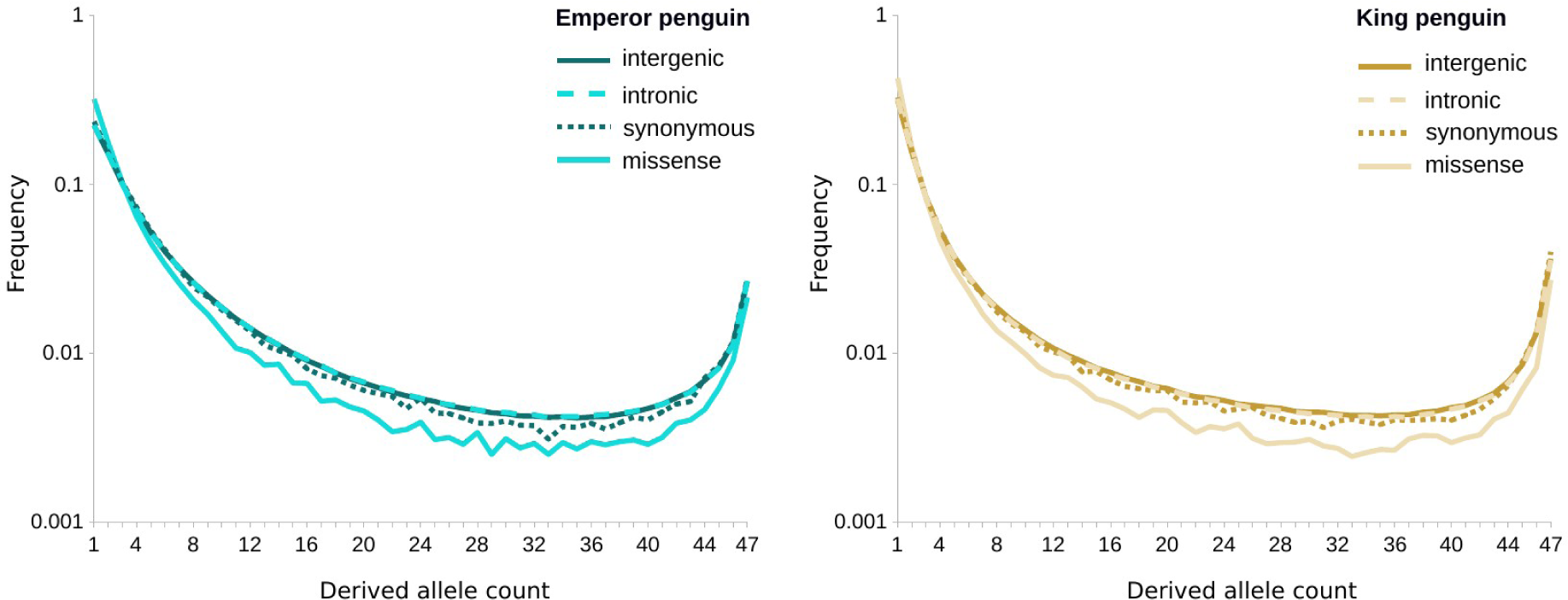
Derived allele frequency distributions for different types of variants (Intergenic, intronic, synonymous and missense as annotated by SNPeff)

#### 1.6 Population-level summary molecular statistics

Nucleotide diversity (*π*) and Tajima’s *D* were estimated in non-overlapping windows of 10 thousand base pairs (kbp) in Emperor and King penguin samples using vcftools across 437 reference genome scaffolds. As we are interested in the species-level long-term adaptation processes, we included in our samples individuals from multiple locations to get an accurate representation of the whole species diversity. Previous studies, based on reduced-representation genome sequencing of hundreds of individuals per species, described both Emperor and King penguin as quasi-panmictic species, with only one very large population each and very little differentiation among colonies (Cristofari et al 2016, 2018). By estimating Weir and Cockerham (1984) *F_ST_* in 10 kbp non-overlapping windows using vcftools (Supp. Fig. 4 inset), we confirmed previous results finding negligible genetic differentiation between King penguin samples from Crozet and Heard, and South Georgia (*F_ST_* mean 0.003, std 0.024) and between Emperor penguin samples from Terre Adélie and Dronning Maud Land (*F_ST_* mean 0.004, std 0.020). Principal component analysis (SNPRelate - Zheng et al 2012, 500 bp pruning for linkage disequilibrium) on a subset of ca. 20,000 SNPs from scaffold NW_008796188 showed no genetic structure in any of the two species (Supp. Fig. 4).

**Supplementary Figure 4.**
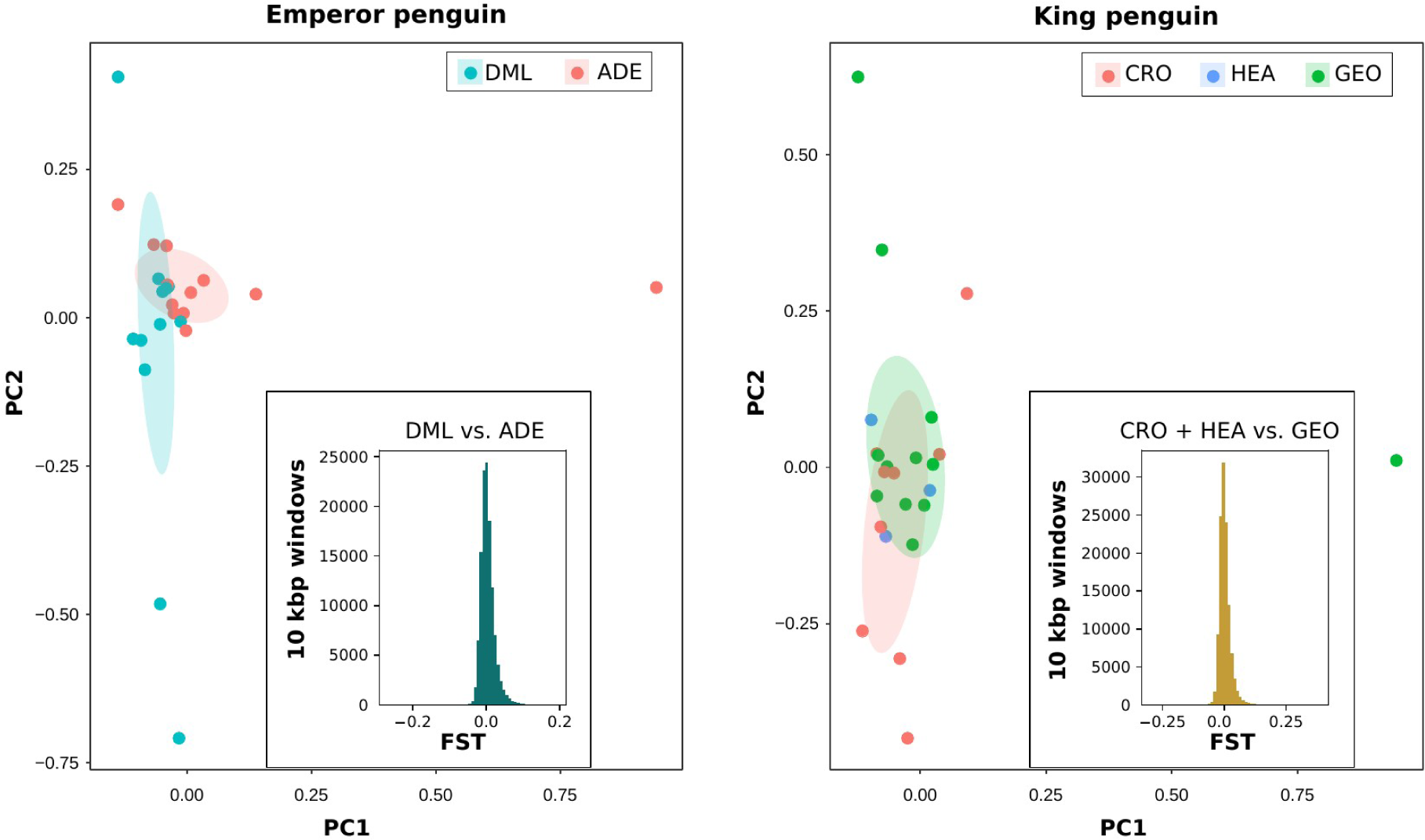
No signature of genetic differentiation among distant colonies of Emperor (left) and King penguin (right). Main plots show the results of Principal Component Analysis run in SNPRelate using 20,256 and 18,256 SNPs not closer than 500 bp from scaffold NW_008796188, respectively. Histograms of *FST* estimated in non-overlapping 10 kbp windows across the whole genome are shown as insets. DML: Dronning Maud Land; ADE: Terre Adélie; CRO: Crozet; HEA: Heard; GEO: South Georgia.

### Gene expression data

#### 2.1 Sample collection and storage

During June-September 2016 field campaigns at Dumont d’Urville Station in Antarctica and at the Alfred Faure station in Crozet archipelago, samples of five different tissues (brain, liver, kidney, skin and muscle) were collected from three freshly-predated 3-7 months-old chicks from natural populations of Emperor and King penguins, respectively (Paris et al 2023). All tissue samples were collected immediately after death, directly fixed in RNAlater (Applied Biosystems, Warrington, UK) and frozen at −80°C until RNA extraction.

#### 2.2 RNA extraction, pooled library preparation and sequencing

Total RNA was isolated from 40 mg of each tissue sample by a standard laboratory-based chloroform extraction after homogenization in 500 ul of TRIzol® reagent (Invitrogen, ThermoFisher Scientific). Samples were added to 100 μl of chloroform, vortexed, and centrifuged at 12,000×g for 15 min at 8°C; the upper aqueous phase was collected and transferred to a new tube for precipitation with isopropanol by centrifugation at 12,000 g for 10 min at 8 °C; the RNA pellet was washed with 75% ethanol and centrifuged at 7,500×g for 5 min at 8 °C, the ethanol removed and the RNA pellet resuspended in RNase free water to be stored at -80C°. As TRIzol-based extraction from skin and muscle yielded poor RNA quality and quantity, likely due to large amount of proteins, connective tissue, and collagen in these tissues, RNA from these two tissues was extracted using the RNeasy Fibrous Tissue Mini Kit (Qiagen) according to the manufacturer’s instructions. Also in this case, isolated RNA was dissolved in RNase free water and stored at -80C°. Concentration and purity (*i.e.,* the A260/A280 ratio) of each RNA sample was assessed by Nanodrop 2000 (Thermo Fisher Scientific) and Qubit 4.0 fluorometer (ThermoFisher Scientific) while RNA integrity was evaluated by capillary electrophoresis on Agilent 2100 Bioanalyzer (Agilent technologies, Santa Clara, CA). As the target of our study was to estimate the global level of gene expression (across tissues), a total of six RNA pools (three pools of five tissues per three individuals for each species) were assembled starting from 15 RNA samples per species, after concentration was normalised. RNA-seq library preparation and sequencing was carried out by BMR Genomics Service (Padova, Italy). Libraries were synthesised using the TruSeq Stranded mRNA Sample Prep kit (Illumina, San Diego, CA), according to the manufacturer’s instructions. Poly-A mRNA was fragmented for 3 minutes at 94°C, and each purification step was carried out with 1 × Agencourt AMPure XP beads. Paired-end sequencing (100 bp from each end) was then performed on the Novaseq 6000 (Illumina, San Diego, CA) at a sequencing depth of 100 million reads per library. Raw reads are publicly available at ENA database as one pool of reads per species (Project ID: PRJEB64484, sample accession ID King penguin: ERS16093259; sample accession ID Emperor penguin, ERS16093260).

#### 2.3 RNA mapping, base-pair and gene expression rate estimates

RNAseq reads from the two penguin species were mapped to the same reference genome (*i.e.*, *A. forsteri* reference genome - RefSeq assembly accession: GCF_000699145.1) as for the genomic data. In particular, after standard filtering and trimming with Trimmomatic, RNA reads were mapped using STAR v.2.7.9a (Dobin et al 2013) and resulting bam files indexed with SAMtools. From bam files, counts of reads overlapping each gene were estimated with HTseq (Anders et al 2015) using the available genes annotation for GCF_000699145.1 reference genome. Multi-mapped and overlapping multiple expression features reads were discarded. For each gene, genomic coordinates, exon and CDS length were extracted from the annotation of GCF_000699145.1 reference genome with the following *bash* commands and merged with the RNAseq expression counts from both Emperor and King penguin samples:

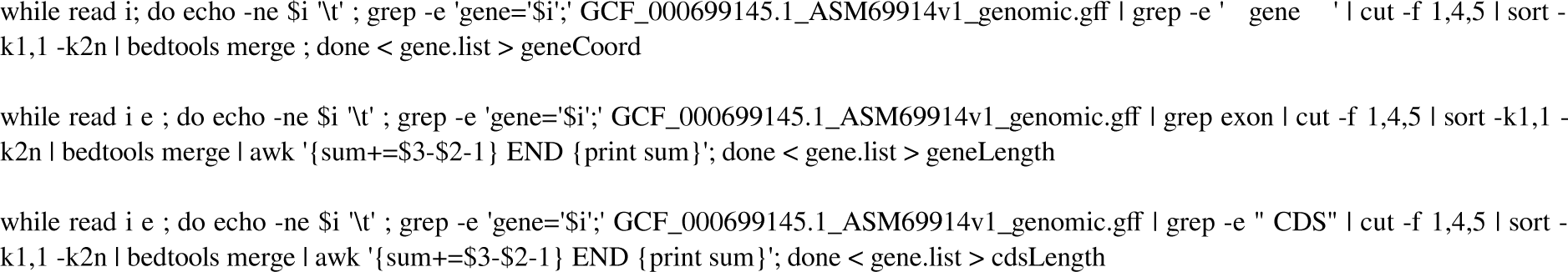

RNAseq counts per gene from HTseq were then normalised by CDS length and total number of reads as transcript per million (TPM), separately per Emperor and King penguin samples (Supp. Fig. 5).

**Supplementary Figure 5.**
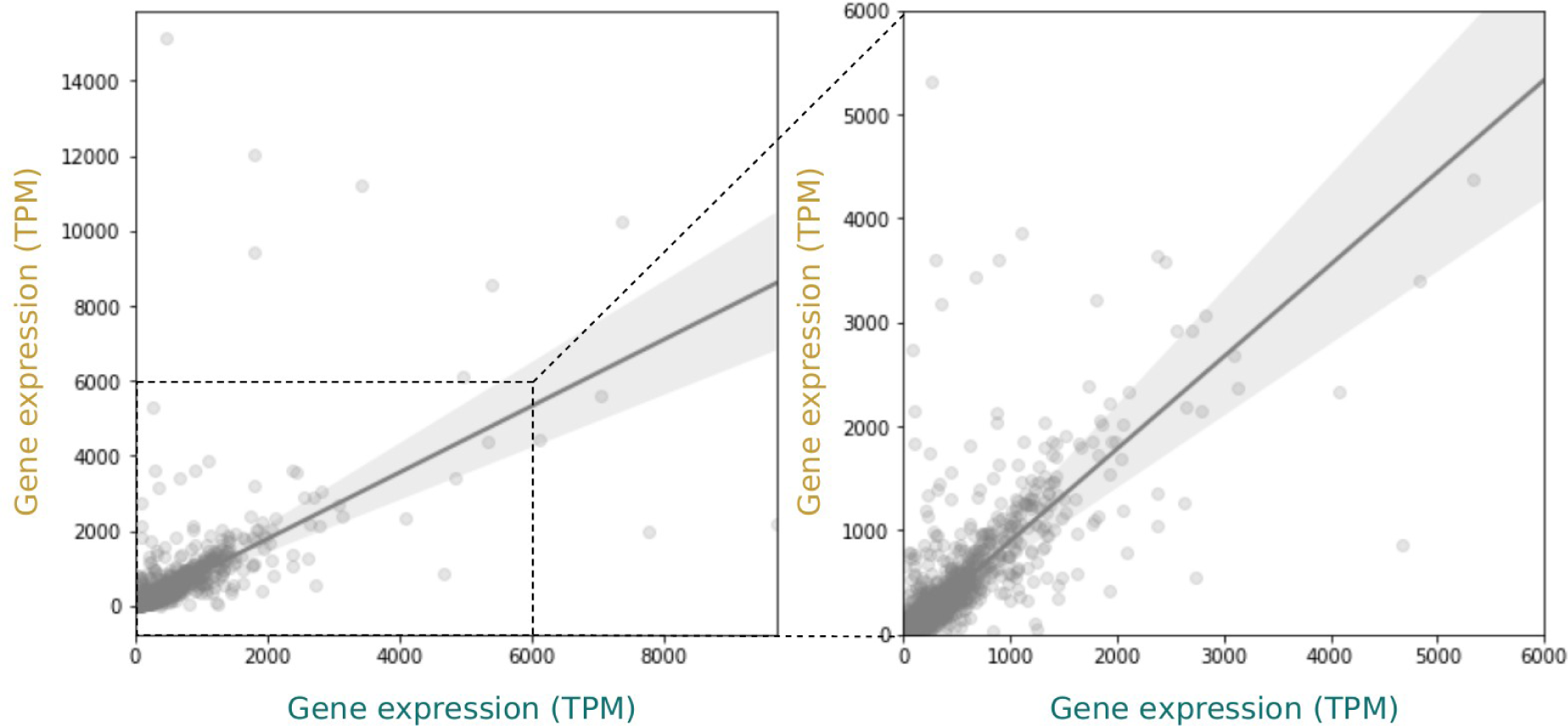
Expression rate as transcripts per million (TPM) per gene in King (gold) and Emperor (teal) penguins. Whole range of expression (left panel) and up to 6000 TPM (right panel).

### Testing the effect of gene expression and population size on purifying selection

#### 3.1 Theoretical expectation

Thermodynamic equations allow us to derive the proportion of protein molecules that are in the native (folded) conformation in the cytoplasm. We assume that each misfolded protein molecule has the same selective cost, caused by its toxicity for the cell. Under this model, the total selective cost of a destabilising mutation is now directly proportional to the total amount of misfolded proteins and it is proportional to the expression level *y*. It is then possible to derive the equilibrium at mutation-selection equilibrium as a function of expression level *y*. Moreover, under different effective population size (*N_e_*), the strength of selection exerted on destabilising mutations is different and thus the equilibrium is different. At the specific equilibrium between mutation, selection and drift, the rate of evolution is given by the probability of fixation of a selected mutation (relative to neutral mutation), called *ω*. In practice, *ω* is approximated from the ratio of nonsynonymous to synonymous polymorphism, *π_N_*/*π_S_*, and or divergence, *d_N_*/*d_S_*. Altogether, it is possible to derive analytically the change in *ω* as a function of *N_e_* and *y* as in eq 18 from Latrille & Lartillot (2021).

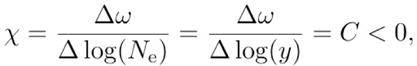

where *C* is a constant that depends on thermodynamic parameters. From this equation, *ω* is linearly decreasing with *N_e_* (in log scale) as well as with *y* (in log scale), importantly the slope of the linear model is the same for both. Additionally, the assumption that proteins are selected against toxicity for the cell can be relaxed and the above equation is also valid more broadly under the assumption that selection is acting on protein-protein interactions (i.e. the protein is bounded or not to other proteins).

**Supplementary Figure 6.**
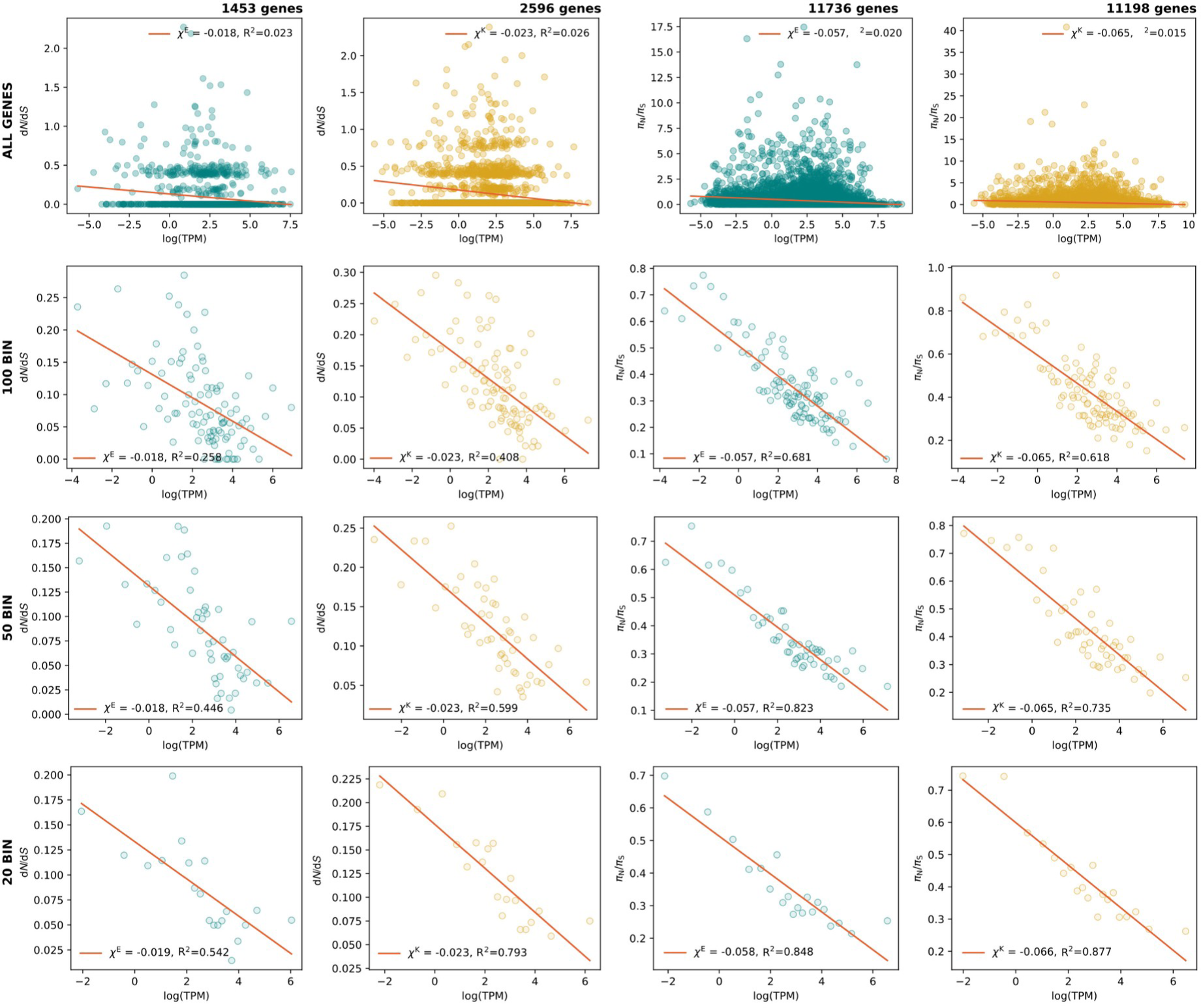
Protein evolutionary rate (*ω*) as *dN*/*dS* (left) or *πN*/*πS* (right) as a function of expression rate (log TPM) in King and Emperor penguins using all genes or binning genes by expression rate in 100, 50, or 20 bins (from top to bottom). *χ^E^* and *χ^K^* are the slopes of the linear regressions for Emperor (teal) and King (gold) penguins respectively. The slope of *πN*/*πS* as a function of log expression level is not dependent on the number of bins used to compute *πN*/*πS* or *dN*/*dS*. However, for fewer bins, the linear model is a strong fit (high *R^2^*), but the fit decreases as the number of bins increases.

#### 3.2 Estimates of synonymous and nonsynonymous polymorphism and divergence per gene

Across different genes, polymorphism and divergence counts are not directly comparable because they are not in the same unit and are mechanically higher for genes with more sites. Moreover, even under neutrality, non-synonymous polymorphism counts are expected to be higher than synonymous counts because a mutation is more likely to be nonsynonymous than synonymous. This argument is also true for nonsynonymous and synonymous substitutions that must be corrected for when estimating species divergence. The number of sites (a.k.a. opportunities of synonymous and nonsynonymous mutations) is thus needed to correct polymorphism and divergence counts and to obtain normalised nonsynonymous divergence (*d_N_*), synonymous divergence (*d_S_*), synonymous polymorphism (*π_N_*) and nonsynonymous polymorphism (*π_S_*) in the same unit.

For each gene, all possible nucleotide mutations were computed from the reference protein-coding DNA sequence (3 x L mutations for a sequence of L nucleotides). Whether a mutation was synonymous or nonsynonymous was determined by comparing the reference codon to the codon obtained after the mutation. Moreover, each mutation was weighted by the instantaneous rate of change between nucleotides, derived from fitting a nucleotide substitution model to the b10k genome alignment (Feng et al 2020).

Counts of synonymous and nonsynonymous polymorphic sites were summarised per gene using a custom python script (*FinalPipeline.ipynb;* https://github.com/emitruc/ExpressionLoad). After applying the stringent filters for coverage (from 6X to 8X per allele) and missing data (no missing genotype), we used the genomic coordinates of all genes to subset the list of SNPs in the *daf.joint.no00* dataset and count the number of synonymous and nonsynonymous polymorphic sites per gene.

The proportion of synonymous mutations was then given as the sum of the instantaneous rates of all synonymous mutations, divided by the sum across all possible mutations (synonymous, nonsynonymous, stop). This proportion of mutations being synonymous is multiplied by the number of sites in the gene to obtain the number of synonymous sites. Repeating this process for nonsynonymous mutations gives the number of non-synonymous sites.

As an estimator of genetic diversity, we used Tajima’s *π*, the average pairwise difference between all sequences in the sample. *π* was obtained for each population from the site-frequency spectrum (SFS) as eq. 5-6 in Achaz (2009).

Formally, *ξ* is a vector that represents the unfolded frequency spectrum composed of *ξ_i_*, the number of polymorphic sites at frequency i/n in the sample (1 ≤ i ≤ n − 1), where n = 48 is the sample size (twice the number of individuals) in the population. π is a function of *ξ* as:

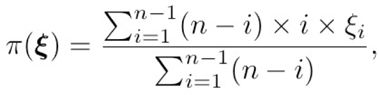

*π* was computed separately for nonsynonymous (*π_N_*) and synonymous (*π_S_*) polymorphism per gene and also globally for intergenic regions (*π_I_*, see below). Finally, to correct *d_N_*, *d_S_*, *π_N_*, *π_S_* such that they are comparable between them, they are expressed in the same unit (per site) by normalising with the number of nonsynonymous or synonymous sites (see above), respectively.

Python scripts used for data handling, parsing, plotting and statistical testing are available at github.com/emitruc/ExpressionLoad and github.com/ThibaultLatrille/PenguinExpression.

#### 3.3 Rate of protein evolution (ω) as function of expression level

*ω* represents the rate of evolution of a protein and it was computed for each gene as the ratio of nonsynonymous to synonymous polymorphism (*π_N_*/*π_S_*) or divergence (*d_N_*/*d_S_*). To compute *π_N_*/*π_S_* or *d_N_*/*d_S_*, each gene must have at least one synonymous count, otherwise the ratio is undefined. *χ* is the slope of the linear regression of *ω* as a function of log(*y*), where *y* is the expression level of the gene in TPM (transcripts per million). We computed *χ* independently in the two penguin populations (Emperor, E) and (King, K), and we denote *χ^E^* and *χ^K^* their estimates of *χ*, respectively (Supp. Fig. 6). To assess the robustness of the results and assess the fit of the linear model, we performed the same analysis while binning genes by their expression level. We performed the analysis with respectively 20, 50 and 100 bins, and computed the slope of the linear regression *χ* and *R^2^* (Supp. Fig. 6).

To compare our results with the patterns shown in Figure 1 of Zhang and Yang (2015), we plot *ω* as *d_N_*/*d_S_* or *π_N_*/*π_S_* in log scale as a function of expression rate (log TPM) in Emperor and King penguins using all genes (Supp. Fig. 7). As *d_N_*/*d_S_* = 0 and *π_N_*/*π_S_*=0 cannot be log-transformed, they are clipped to *d_N_*/*d_S_* = 10e-3 and *π_N_*/*π_S_* = 10e-3, respectively, as in Zhang and Yang (2015). The pattern shown by *π_N_*/*π_S_* is similar to those presented in Figure 1 of Zhang and Yang (2015), in particular in the case of *Drosophila melanogaster*, *Mus musculus* and *Homo sapiens*. The very recent divergence between the two penguin species resulted in many genes with *d_N_*/*d_S_*=0.

**Supplementary Figure 7.**
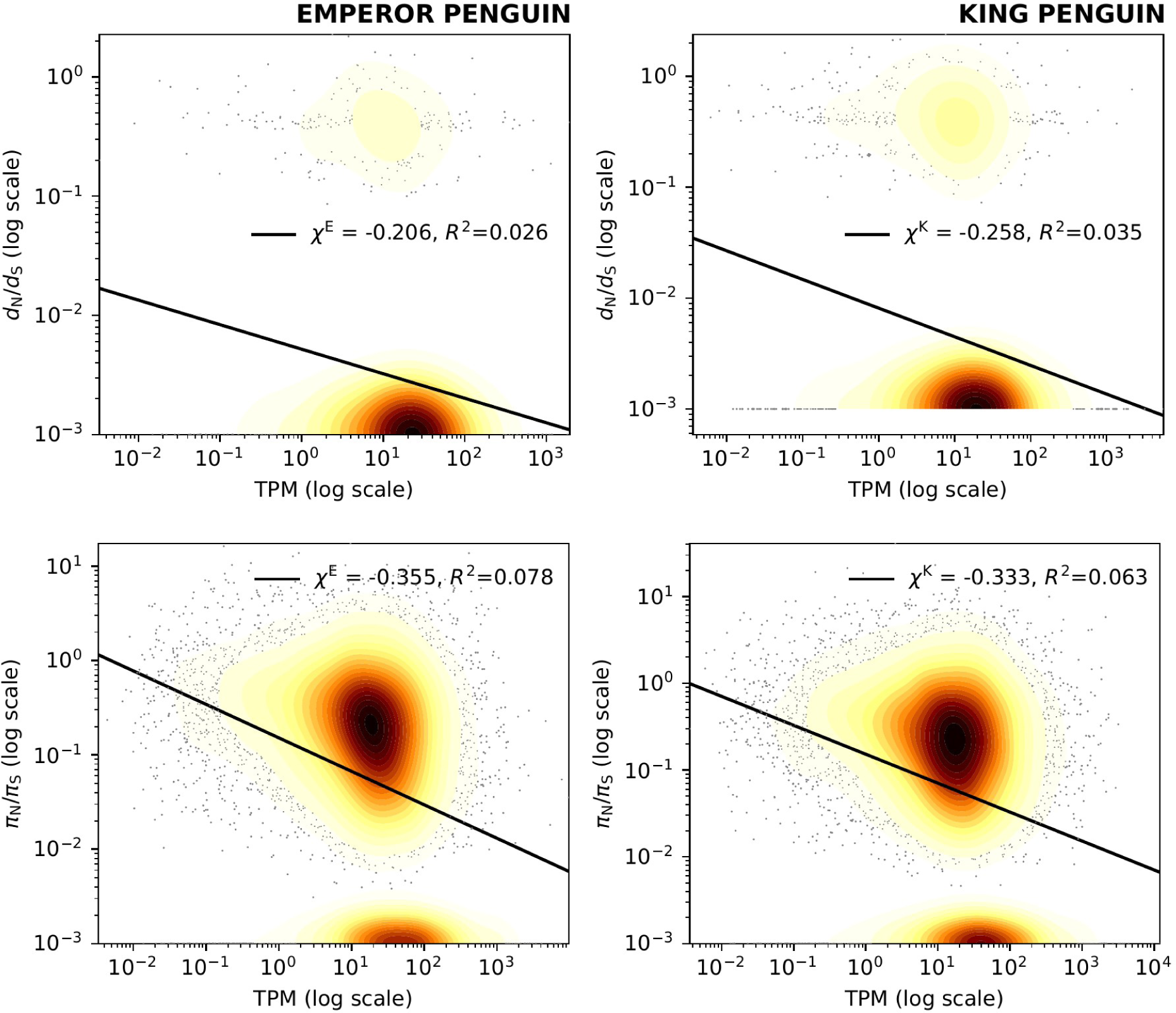
Protein evolutionary rate (*ω*) as *dN*/*dS* (top) or *πN*/*πS* (bottom) in log scale as a function of expression rate (log TPM) in Emperor penguin (left) and King penguin (right) using all genes.

#### 3.4 Rate of protein evolution (ω) as function of effective population size (N_e_)

Given the two penguins populations (Emperor, E) and (King, K), we also estimated *χ* as the change of *ω* as a function of *N_e_*:

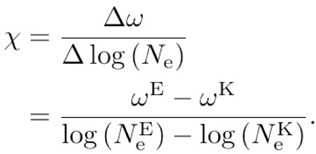

Under the assumption that the mutation rate (*u*) is the same between the two species, and since *π* = 4 *N_e_ u* from neutral markers, *χ* simplifies to:

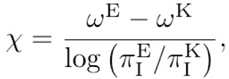

where *π ^E^*and *π_I_ ^K^* are estimated from the intergenic regions, which are assumed to be neutral. Here normalisation by the number of sites is not required since *π^E^* and *π^K^* are already expressed in the same unit in the two species (the same reference genome was used), which cancels out in the ratio *π_I_ ^E^*/*π_I_ ^K^*. *ω* (either *π_N_*/*π_S_* or *d_N_*/*d_S_*) is computed as the total count (polymorphism or divergence) across all genes, divided by the total number of sites across all genes, respectively for polymorphism and divergence. We performed a bootstrap sampling (1000 replicates) to estimate the confidence interval of *χ*, where genes were sampled with replacement in each replicate (Supp. Fig. 8).

**Supplementary Figure 8.**
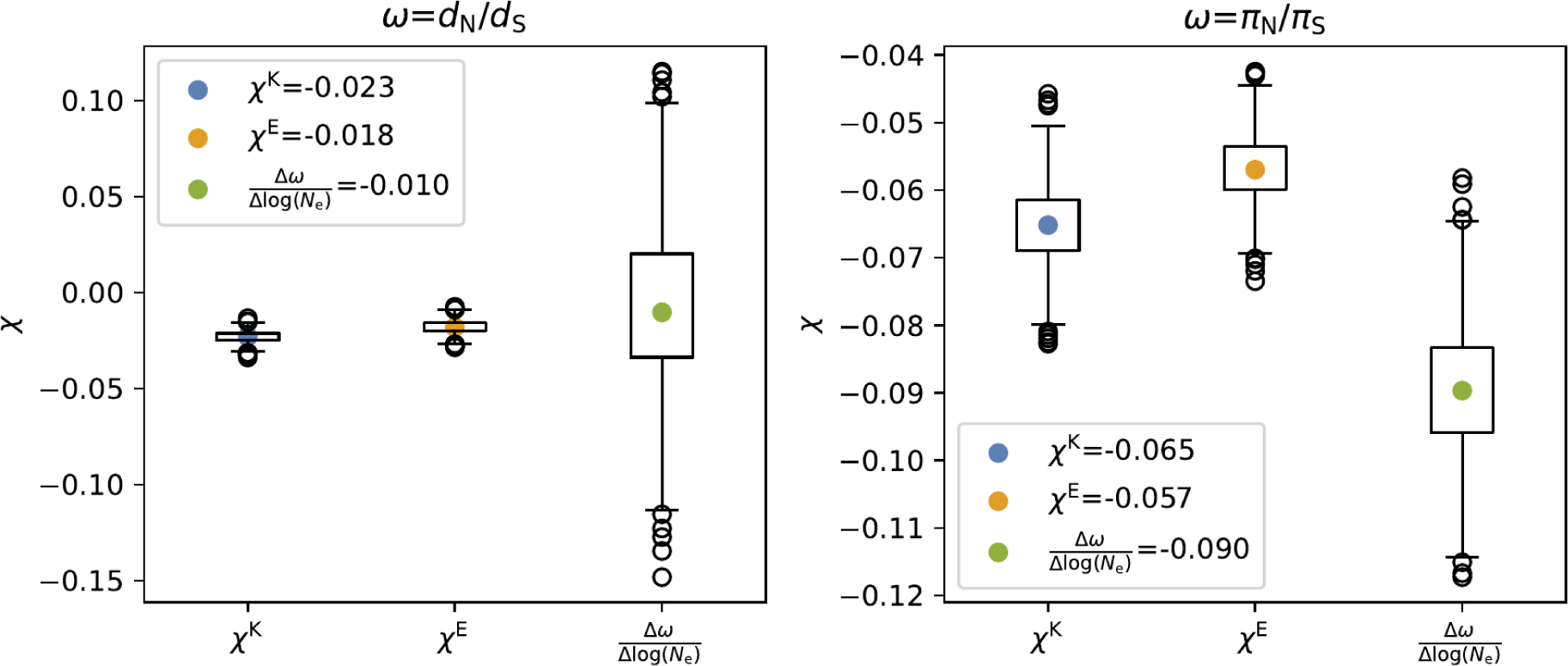
Rate of protein evolution (*ω*) as function of expression level and of effective population size (*Ne*). *χ* is the slope of the linear regression of *ω* (either *dN/dS* in left panel *πN/πS* in right panel) as a function of the expression level of the gene in TPM (transcripts per million) in the log scale. We computed *χ* independently in the two penguin populations (Emperor, E) and (King, K), and we denote *χ^E^* and *χ^K^*their estimates of *χ*, respectively. *χ* is also estimated (third column) as the change of *ω* as a function of *Ne* (section 3.4). Confidence intervals of *χ* are obtained from bootstrap sampling (1000 replicates) where genes are sampled with replacement in each replicate.

#### 3.5 Polymorphic sites-based analyses

For each polymorphic SNP in our genomic dataset (*daf.joint.no00*), we separately estimated the RNA reads coverage in King and Emperor penguin samples using SAMtools module *depth* and a bed file listing all polymorphic sites in the genomic dataset. Per site RNA read coverage was then added to the genomic dataset, as one value (total number of reads) per species (*daf.joint.no00.rnaCov;* https://doi.org/10.6084/m9.figshare.23863503.v1). Raw mRNA coverage per site was normalised as counts per million reads (CPM) dividing this value by the sum of mapped RNA reads x 1 million, separately per species:

emp_counts = [62589631 + 85819833 + 72587896] #mapped reads in each of the three pools
king_counts = [66803162 + 64732729 + 58348853] #mapped reads in each of the three pools

emp_CPM = emp_RnaCov / emp_counts * 1000000
king_CPM = king_RnaCov / king_counts * 1000000

After applying the same stringent filters for coverage (from 6X to 8X per allele), outgroup missing data (at least 4 alleles present), and King and Emperor penguin missing data (no missing genotype allowed), we generated one dataset of synonymous and nonsynonymous variants together per species. Next, after capping the maximum CPM value to 5, we grouped the variants in each dataset applying 100 bins of CPM. We then estimated the ratio of nonsynonymous over synonymous variants in each bin separately per species (Supp. Fig. 9). To investigate the effect of gene expression on the allele frequency of synonymous and nonsynonymous variants, we estimated the derived allele frequency spectra grouping the variants by discrete values of CPM (Supp. Fig. 10): *i*) CPM < 0.3, CPM > 0.3; *ii*) CPM < 0.5, CPM > 0.5; *iii*) CPM < 0.3, 0.3 < CPM < 2, CPM > 2. In order to exclude the possibility that a few genes were driving the observed pattern (*i.e.*, pseudoreplication) in the comparison of the site frequency spectra at different expression rates, we replicated ten times the analysis by grouping variants with CPM < 0.3 or CPM > 0.3 after randomly subsampling one synonymous and one nonsynonymous variant per gene. Out of ten replicas, we estimated the 95% intervals for each derived allele count in the site frequency spectra (Figure 3 in main text).

**Supplementary Figure 9.**
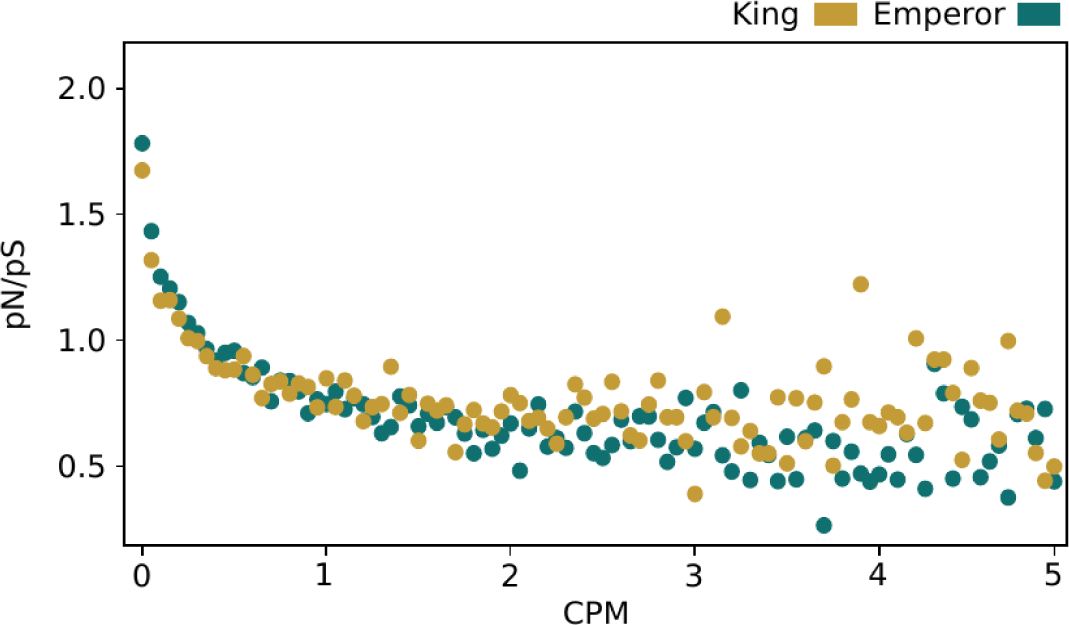
Ratio of missense to synonymous segregating sites per 0.05 intervals of counts per million (CPM) mRNA coverage of each site. Total mRNA read coverage (normalised as counts per million reads) across five tissues from three specimens has been scored for all nonsynonymous and synonymous sites. Note these are density histograms where the total sums up to 1 in each species.

**Supplementary Figure 10.**
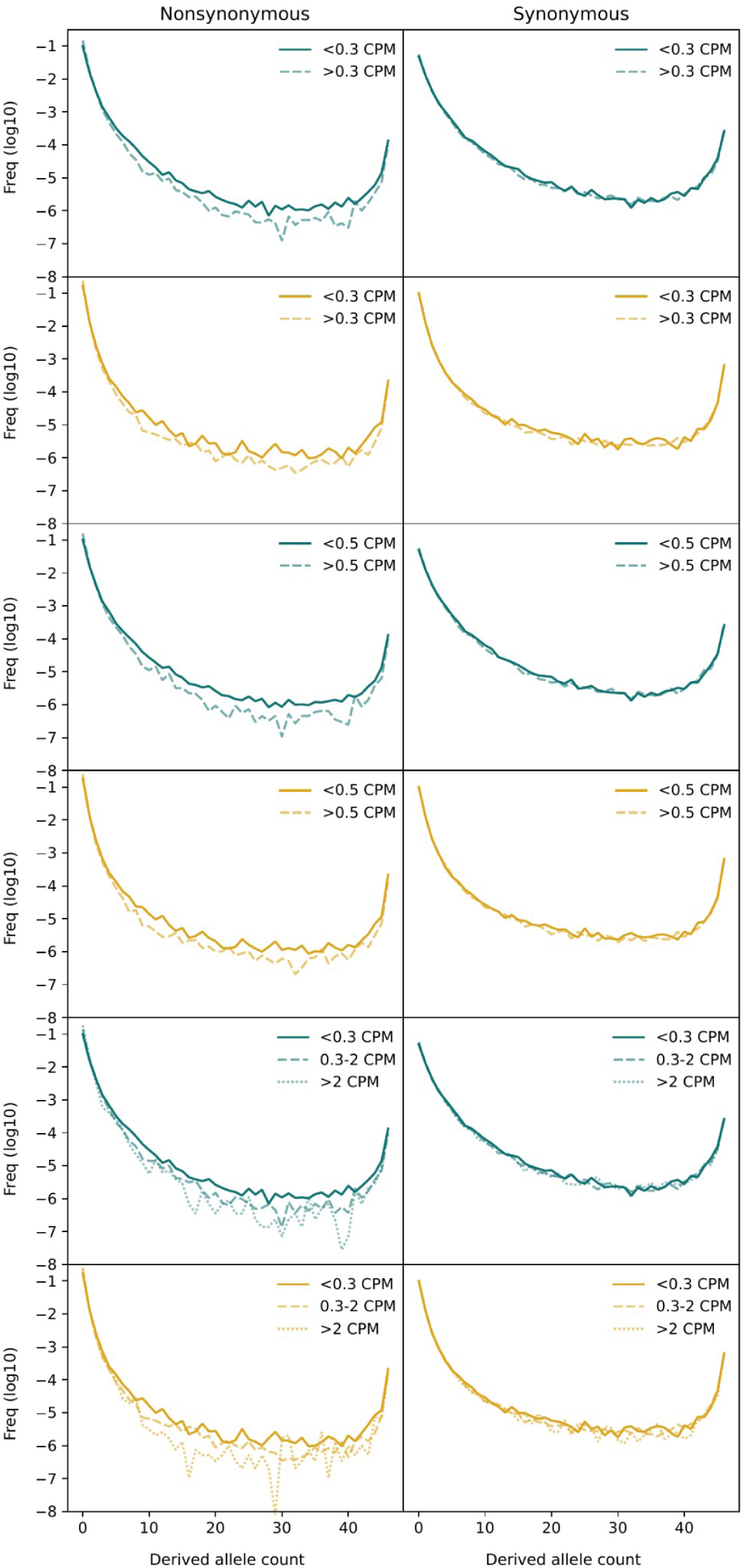
Site frequency spectra of all nonsynonymous (left column) and synonymous variants (right column) across all genes of Emperor (teal) and King (gold) penguins with mRNA expression partitioned by different values of CPM. The relative frequency of each count class is log10 transformed.

### Estimating the selection coefficients of highly expressed genes using realistic forward simulations

The genomic and transcriptomic data analysed in the previous section show that *i)* purifying selection on genes appear as correlated with their level of expression, and that *ii)* the effect of gene expression on purifying selection overrides the effect of population size at highly expressed genes. To infer the selection coefficients causing the pattern of purifying selection observed at highly expressed genes, we devised a forward in time genomic simulation framework in SLiM v4.0.1 (Haller and Messer 2023). In particular, we used both general Wright-Fisher (WF) and penguin-specific non Wright-Fisher (nonWF) models to study the effect of different population sizes (*i.e.*, genetic drift) and the effect of different selection coefficients on *p_N_*/*p_S_* (which is the same as *π_N_*/*π_S_* when analysing simulated data). Our hypothesis was that the effect of demography (*N_e_*) alone isn’t strong enough to generate the *p_N_*/*p_S_* values as observed in the King and Emperor penguin data, but much stronger selection coefficients are needed in the model to explain the pattern observed at highly expressed genes.

Firstly, to investigate the correlation between population size (*N_e_*) and *p_N_*/*p_S_*, we designed a WF (“*WF Pop Size Effect Model”*) and a nonWF model (“*nonWF Pop Size Effect Model*”), which we ran with different carrying capacities (*N_e_* = 1,000, 10,000 and 100,000). The main difference in the nonWF model is that we modelled realistic King penguin life history traits (Céline Le Bohec, *personal communication*) with population size as a non-fixed parameter which can fluctuate around the set carrying capacity. The genomic model is the same in WF and nonWF simulations and it is implemented as a reduced version of the whole CDS of the King penguin with the coding component of 1,000 genes of length 2,400 bp (*i.e.*, the mean value for the coding sequence per gene in King penguin). The recombination rate is set at 1e-8 within genes and at 4.8e-4 between genes (1e-8 rate scaled to the average intergene length). Mutation rate is set at 1e-8 and the ratio between the occurrence of deleterious and neutral mutations is 2.31:1 (Kim et al 2017). The selection coefficient is assigned to each deleterious mutation using a random value from a gamma distribution with mean - 0.01314833 and shape 0.186 (Kim et al 2017). This distribution has been used before in humans and other mammals but we believe it could approximate the distribution of fitness effect also in our target species. For the dominance coefficient we used a *h*-mix model (Kyriazis et al 2021), where weakly deleterious mutations (*s* ≥ −0.01) are partially recessive (*h* = 0.25), while strongly deleterious mutations (*s* < -0.01) are totally recessive (*h* = 0).

We designed similar WF and nonWF models (“*WF Gene Expression Effect Model*” and “*nonWF Gene Expression Effect Model*”, respectively) to investigate the effect of gene expression on *p_N_*/*p_S_*. As the *p_N_*/*p_S_* in the largest simulated population size (*N_e_* = 100,000) of the *Pop Size Effect* models did not reach small values as observed in our King and Emperor penguin data, we implemented a more extreme selection scenario. In the initial *Pop Size Effect* models, the gamma distribution used to randomly assign the selection coefficient of a novel deleterious mutation resulted in most of the mutations being weakly deleterious (*s* >= -0.01). Here, we assigned a fixed selection coefficient (*s* = -0.001, -0.01, and -0.1) to every gene so that all nonsynonymous mutations appearing in a gene have the same selection coefficient, then we simulated 100 genes for each selection coefficient resulting in a set of 300 genes. We followed the same strategy for the dominance coefficient assignment, thus it is assigned to every gene deriving it from its fixed selection coefficient following the *hs* relationship (Kyriazis et al 2021), so that dominance and selection coefficients result to be inversely proportional:

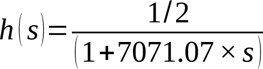

The recombination coefficient is set at 1e-8 within genes and 0.5 between genes in order to make genes independent to one another. Since most of the genes resulted in a low number of mutations at the end of the simulated generations and to keep the computational running time tractable, we instead increased the gene length from 2,400 to 34,000 bp (the average total gene length in King penguin genome) in the final models, we then estimated the *p_N_*/*p_S_* per each selection coefficient category of genes under different carrying capacities (*N_e_* = 1,000, 10,000, 100,000).

In total, we designed four models, each of them testing three carrying capacities. For each of these 12 scenarios, we ran three replicas of 10\**N* generations each (except for the *N_e_* = 100,000 model where we ran for *N_e_* generations due to computational time), as suggested by SLiM authors in order to make the population reach an equilibrium state (*i.e.*, burn-in). At the end of the simulations, we estimated the *p_N_*/*p_S_* based on 24 individuals, as in our genomic and transcriptomic real data, repeating the estimate 100 times by randomly resampling 24 individuals (Supp. Fig. 11). Slim scripts are available at github.com/PiergiorgioMassa/penguin_gene_expression_slimulations.

**Supplementary Figure 11.**
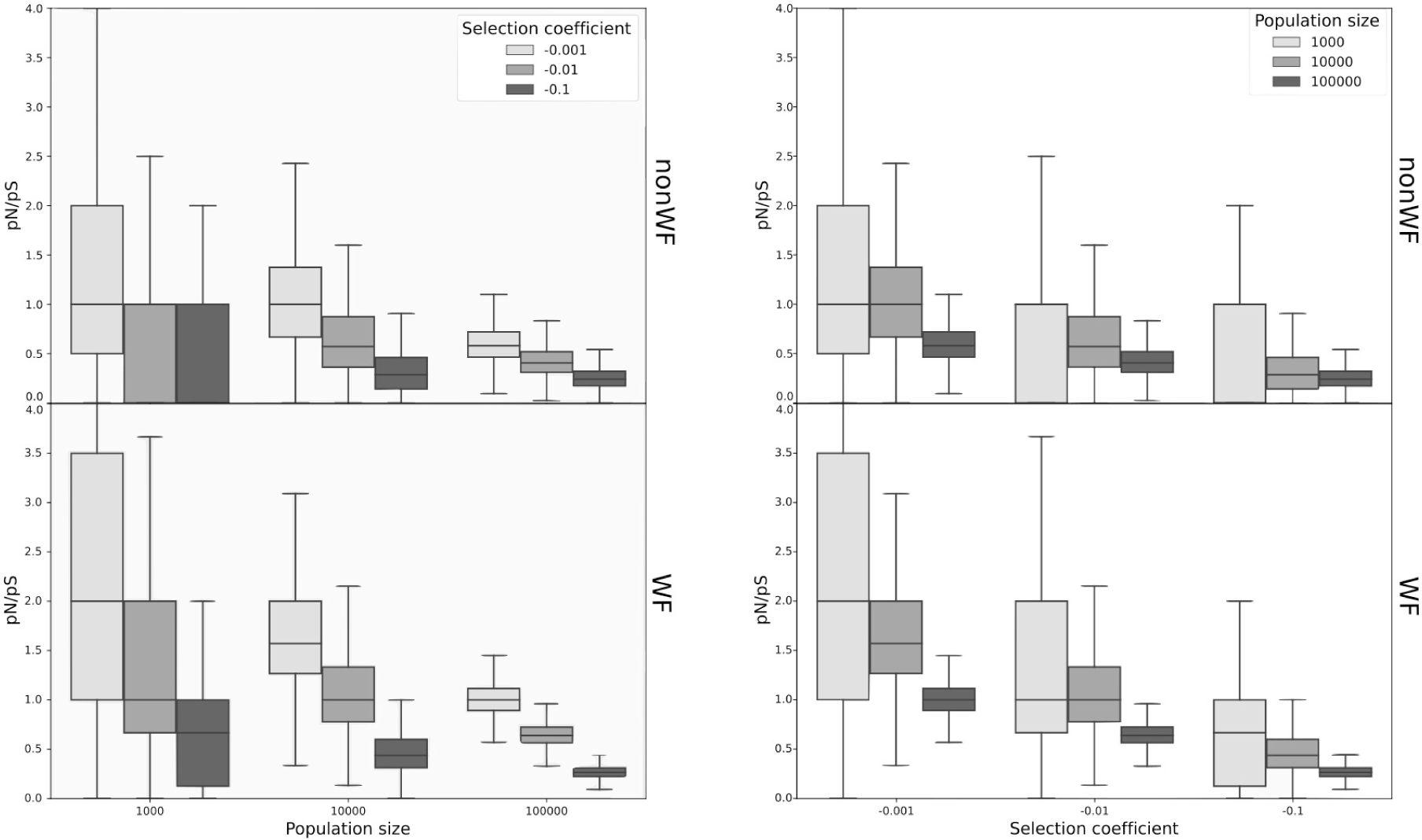
Selection coefficients in highly expressed genes. Distributions of *pN*/*pS* estimated in 100 genes where each deleterious mutation is assigned a fixed selection coefficient of -0.001, -0.01, or -0.1 in simulated populations of 1,000, 10,000, or 100,000 individuals, plotted by population size (left panels) or selection coefficient (right panels). As expected, stronger selection coefficients (i.e., *s* = -0.1) are effective also when population size is small.

### Expression rate per fitness effect class predicted by SNPeff

Genetic load is the cost paid by any population to its potential further evolution. It is an inherent feature of populations evolving by random mutations which more often have deleterious than advantageous effects on fitness. Deleterious mutations can accumulate in some individuals as a consequence of small population size (i.e., high genetic drift and high inbreeding) reducing the fitness of these high load individuals as compared to individuals which bear fewer such mutations. In conservation genomics, genetic load is getting growing attention as a more appropriate measure of a population’s genomic health (Bertorelle et al 2022). However, genetic load is difficult to estimate in non-model species, especially when relying on genomic data only, without information on mutations’ fitness effect. In such a case, genetic load can be estimated using *i)* either the predicted effect of a mutation on the amino acid sequence (Cingolani et al 2012), *ii)* or the evolutionary conservation of a certain allele at orthologous sites across multiple species (*i.e.*, GERP; Davydov et al 2010). The latter has the advantage that it can be applied even outside coding regions, but on the other hand, it requires large multi-species genomic alignments, which are extremely computation intensive and error-prone. Even if it can only be applied to coding sequences, a genomic load proxy based on gene expression could be more accurate and easier to compute, given that mRNA expression data from multiple tissues of the target species (or one closely related) are available.

After excluding intergenic variants from the *daf.joint.no00.rnaCov* file with a simple bash command (*awk ’$14 != “intergenic”’ daf.joint.no00.rnaCov > daf.joint.no00.rnaCov.noIntergenic*) and applying the same filters for coverage (between 6X and 8X per allele) and missing data in ingroup (no missing) and outgroup samples (at least 4 alleles present for polarisation) as before, we applied an additional Z-normalisation to the CPM for clarity of interpretation only.

#CPM normalization
eCPMscal = sum of rnaCov_emp / 1000000
kCPMscal = sum of rnaCov_king /1 000000
eCPM = rnaCov_emp / eCPMscal
kCPM = rnaCov_king / kCPMscal

#CPM Z-normalisation
eCPMstdz = eCPM - eCPM_mean / eCPM_std
kCPMstdz = kCPM - kCPM_mean / kCPM_std

Next we selected the variants based on their fitness effect predicted by SNPeff (Cingolani et al 2012): LOW-synonymous (we excluded all of the LOW which are present in introns, mostly as splice-region-variants, as these are not covered by mature mRNA seq data), MODERATE, and HIGH effect.

We then summarised the Z-normalised CPM mean and standard deviation across all private variants (segregating and fixed) or only in private fixed variants in each species per SNP effect (Table 1 in the main text).

As site-specific expression of HIGH deleterious variants (mainly start/stop codons loss/gain and splice acceptor/donor variants) could be biassed in mature mRNA sequencing due to their sequence position, we also estimated their expression using the expression rate of the gene they occurred in. Using the gene-based expression data generated before, we tested whether the expression rate of the group of genes with HIGH deleterious derived alleles was significantly lower than all the rest of the genes. Even if we applied a rather conservative test, the expression of genes with predicted highly deleterious variants is on average three times lower than all genes (Kolmogorov-Smirnov test *p-*value = 0.00095) in the King penguin and slightly lower, even if not significant, in the Emperor penguin.

MODERATE variants show the same average expression rate in both Emperor and King penguin samples (0.78) while a higher average expression in private fixed variants, with a larger difference in Emperor (1.41) than King (0.93) penguin sample. The variance in expression is also larger in fixed than in segregating variants (Table 1 in the main text). Instead of a major role of purging, which could anyway still be part of the process, derived nonsynonymous (which mainly contribute to the MODERATE effect variants) alleles which are fixed at highly expressed genes could actually be truly advantageous variants fixed by positive selection. The lower average expression in King penguin is in line with the expected larger effect of random drift in this population leading to fixation of a larger number of derived nonsynonymous variants (2229 in the King penguin as compared to 1166 in the Emperor penguin), but in genes with lower expression rate and, hence, lower effect on individual fitness. More purging could be again suggested in this case, limiting the fixation of deleterious missense in highly expressed genes. We also suggest that the more intense purging of deleterious variants in the King penguin could be the cause of the lower average expression of synonymous variants which were reduced in more expressed genes by background selection (Table 1 in the main text).

To test our hypothesis that private fixed derived MODERATE variants could actually be truly advantageous, we screened the Emperor and King penguin genome data for selection signatures using Sweepfinder2 (De Giorgio et al 2016) and OmegaPlus (Alachiotis et al 2012) in windows of 10 kb, following the authors’ instructions with default settings. Fixed derived alleles are in regions showing higher signatures of selection and lower diversity in both species, and higher differentiation between the two species, hallmarks of highly conserved and lowly recombining genomic regions (Supp. Tab. 3). This has to be considered as a preliminary indication of the potentially positive effect of some of the private fixed MODERATE variants in each species. More targeted analyses are, however, necessary to conclude on the fitness effect of fixed differences with a MODERATE effect on fitness.

**Supplementary Table 3.**
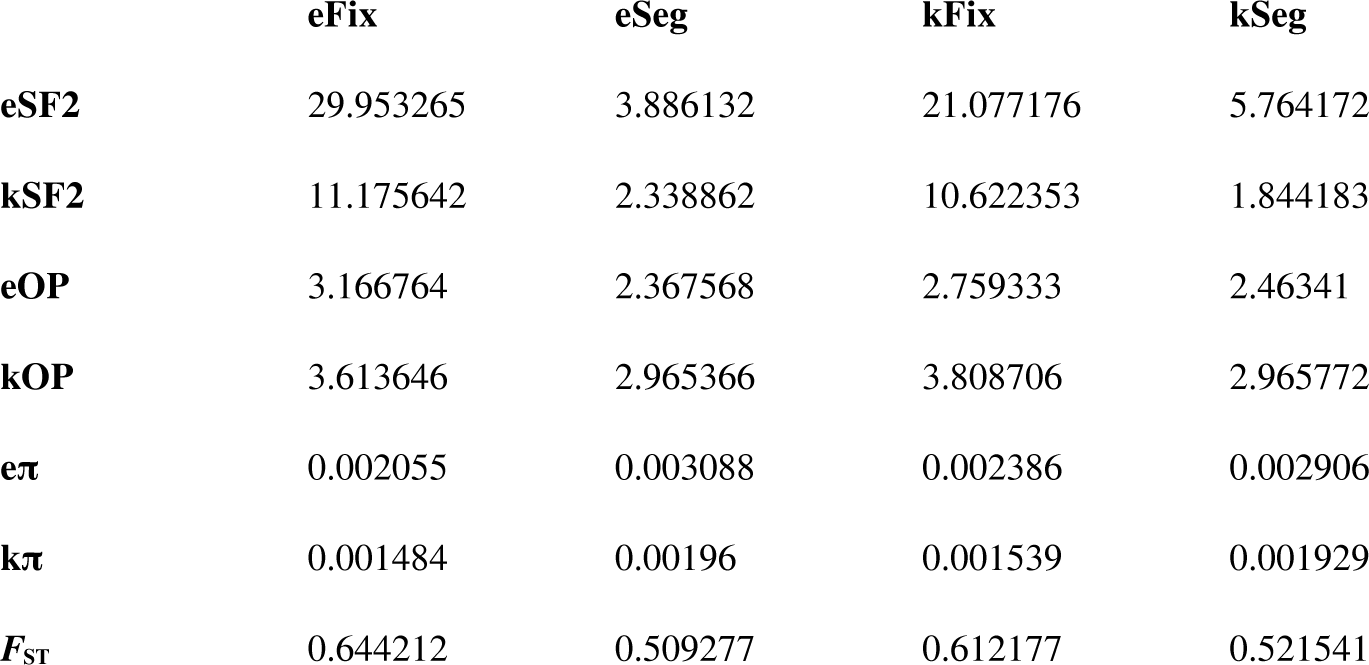
Signatures of selection (SF2: Sweepfinder2; OP: Omega+), nucleotide diversity (π) and differentiation between King (k) and Emperor (e) penguins (*F_ST_*) in 10 kb genomic regions with fixed differences (Fix) or segregating variants (Seg) identified of MODERATE effect by SNPeff annotation. All comparisons between the distribution of the statistics across regions with fixed differences and segregating variants for both species are significant (Kolmogorov-Smirnov test *p-*value < 0.05).

|  | eFix | eSeg | kFix | kSeg |
| --- | --- | --- | --- | --- |
| **eSF2** | 29.953265 | 3.886132 | 21.077176 | 5.764172 |
| **kSF2** | 11.175642 | 2.338862 | 10.622353 | 1.844183 |
| **eOP** | 3.166764 | 2.367568 | 2.759333 | 2.46341 |
| **kOP** | 3.613646 | 2.965366 | 3.808706 | 2.965772 |
| **eπ** | 0.002055 | 0.003088 | 0.002386 | 0.002906 |
| **kπ** | 0.001484 | 0.00196 | 0.001539 | 0.001929 |
| **$F_{ST}$** | 0.644212 | 0.509277 | 0.612177 | 0.521541 |

## Notes

https://doi.org/10.6084/m9.figshare.23863503.v1

https://github.com/emitruc/ExpressionLoad

https://github.com/PiergiorgioMassa/penguin_gene_expression_slimulations

https://github.com/ThibaultLatrille/PenguinExpression

https://zenodo.org/doi/10.5281/zenodo.10688853

### Competing Interest Statement

The authors have declared no competing interest.

### Summary of Updates

The manuscript has been revised after a first round of peer review in Peer Community in Evolutionary Biology. The Recommender and 3 reviewers suggested a few modifications which are fully detailed in the response that will be submitted to PCI Evol Biol and published alongside the article once it will be acceptable and recommended in PCI.

